# A framework for identifying transcript orthologs: the evolution of sex bias in alternative transcript structure in Drosophila

**DOI:** 10.64898/2026.05.25.727716

**Authors:** Kinfeosioluwa S. Bankole, Lauren M. McIntyre, Michael Garan, Alison M. Morse, Netanya Keil, Ashlee Hernandez, Olga Barmina, Mursalin Khan, Artyom Kopp, Rebekah L. Rogers, Rita M. Graze

**Author notes:** co-first authors (Kinfeosioluwa S. Bankole, Lauren M. McIntyre). co-corresponding authors: **Error! Hyperlink reference not valid.**.

## Abstract

**Background:** Recent advances in long read technologies provide an unprecedented opportunity to study transcript evolution. However, comparative evolutionary studies, even in Drosophila, are limited by inconsistent and incomplete annotation, and the lack of annotated transcript homology.

**Results:** In this study of five species spanning 28 million years (*D. melanogaster*, *D. simulans*, *D. yakuba*, *D. santomea* and *D. serrata*), we infer transcript homology using reciprocal liftover, and orthology using network analyses, with data validation from long read RNA-seq of male and female head tissue. We build the first genus level annotation, with 15,996 genes and 56,370 transcripts. Expressed transcripts are conserved, 73% of transcript orthologs are detected in all species. Even the improved annotation underestimates the number of genes with alternative transcripts, with 75% of genes expressing multiple structurally diverse transcripts. In a replicated quantitative evaluation of ∼10,000 genes, both male and female-biased transcripts are expressed in 410 (*D. melanogaster*), 608 (*D. simulans*), and 493 (*D. serrata*) genes and in 118 orthologous genes in the *D. melanogaster* - *D. simulans* species pair, indicating greater potential for resolution of sexual conflict by alternative transcription than previously appreciated. We identified 605 transcript orthologs conserved for sex bias in the *D. melanogaster-D. simulans* species pair and of these, 22 male and 19 female-biased transcripts were conserved in sex bias with the outgroup *D. serrata,* including transcripts of genes involved in brain development, Sxl target *Glutamine synthetase 2* and *ciboulot*.

**Conclusions:** Conserved alternative transcripts suggest that transcriptional diversity is a pervasive driver of the evolution of functional diversity.

## Background

Alternative transcription and alternative splicing are fundamental mechanisms of eukaryotic gene regulation. Altering the 5’ or 3’ exon positions or length, and shuffling combinations of exons, allow a single gene to produce multiple transcripts that can be distinctly regulated and often encode different gene products. For example, recent and ongoing efforts to annotate the human genome find approximately 20,000 protein coding genes corresponding to about 10 times as many annotated transcripts^1^. Such transcriptional diversity underlies tissue and cell specific function, as well as differences between sexes and developmental timepoints. Understanding how transcript diversity evolves, and the extent to which transcripts are conserved, can provide insight into mechanisms underlying the evolution of genome function^2, 3^. Drosophila as a genus has many annotated genomes and is an important model for understanding how the evolution of transcript diversity affects gene function. For example, the presence of both male and female specific transcripts of *Sex lethal* (*Sxl*) underlie a switch that triggers a conserved alternative splicing cascade resulting in somatic sex determination (reviewed in^4^). In Drosophila, there are approximately 14,000 protein coding genes, with a little over two times as many currently annotated transcripts^5^, with the majority of the genes having only a single annotated transcript. How much transcript diversity is conserved across species is unknown. This is due in part to differences in the quality of transcript annotation across species, presenting a challenge to investigating comparative evolutionary hypotheses.

One hallmark of diversity in gene function is context dependent expression^6^. mRNA transcript isoforms of multi-transcript genes may vary in expression by tissue/cell-type, sex, developmental stage and environment^7–10^. While expression variation is insufficient to prove sub or neo-functionalization^11^, conserved context dependent structural variation in transcript use suggests a potential link to function. For example, the known conservation of functionally relevant structural differences in transcripts between sexes^12–14^ and tissues/cell-types^15, 16^. A comparative approach to transcript evolution allows us to generate hypotheses about potential links between diversity in transcript structure and the diversification of function and to better understand the evolutionary context in which new transcripts evolve. There have been many studies of comparative evolution of splicing using short reads, typically by cross species mapping to the most well annotated genome. Despite ongoing efforts to improve analytical power, there are inherent limitations in the accuracy of transcript reconstruction from short reads resulting in an accumulation of biases due to mapping reads across species and/or variation in genome and annotation quality^17–19^.

Long read sequencing technologies have, for the first time, made possible comparative transcriptomic studies based on direct observation of the sequence of complete transcripts. The whole transcriptome of any species for which tissue can be procured can be sequenced, a revolution in our ability to study and understand the evolution of transcription. This has resulted in a more comprehensive view of transcribed regions of genomes^18^. The technology is now affordable enough to use long reads to quantify and statistically compare expression across conditions^20^. Comparing expression patterns between species requires the identification of the ‘same’ transcript in different species, the orthologous transcript. Although there is currently no agreed upon definition of a transcript ortholog, progress has been made in the area^21–24^. Here we develop a systematic framework linking homologous transcripts between species for the whole transcriptome - and in many instances, establish a one-to-one relationship among homologous transcripts in all five species that we use to infer likely orthology.

Gene expression, including transcription and RNA processing, is complex and inherently noisy, with transcript structure and abundance impacted by stochastic elements of regulatory mechanisms^25, 26^. While long reads provide increased resolution, ongoing technological challenges include fragmentary and partial transcripts, combined with relatively large amounts of starting material needed for high quality full transcriptome coverage^18, 27^. Identifying likely transcript fragments depends upon high quality genome annotations^28, 29^, underscoring the need for high quality annotations in multiple species in comparative transcriptomic experiments.

We use high coverage long read RNA-seq in a comprehensive study of male and female head tissue in two pairs of sister species, *D. melanogaster* and *D. simulans*, and *D. yakuba* and *D. santomea,* with *D. serrata* as an outgroup. These species pairs are members of a well-studied comparative evolutionary biology model clade, with a focus on genetic differences in pigmentation, mating behavior, and reproductive isolation^30–35^ (Supplementary Figure 1). *D. serrata* was selected as an outgroup because of the availability of genomic data and detailed studies of sexual selection^36, 37^. For *D. melanogaster*, *D. simulans* and *D. serrata* we replicated long read RNA-seq at high coverage, enabling the first whole transcriptome quantitative comparison of the conservation and divergence of full transcripts and their structure, as well as alternative transcript use between sexes.

Our study presents a roadmap for the successful analysis of the typical scenario in comparative transcriptomic studies, where one species (in this case *D. melanogaster*) has a high-quality transcriptome and genome and the other species are more variable in both the quality of the genome assemblies and the annotations. Using a multilayer reciprocal liftover approach, we build a comprehensive annotation for the genus (the fiveSpecies annotation) that expands the current annotation in all species and links homologous transcripts across all five genomes. We use the empirical data to validate ‘lifted’ annotations and to resolve ambiguity. The resulting robust set of annotations can be widely used by the Drosophila community to evaluate conservation and divergence in these species, and the approach can be extended in a straightforward manner to include more species in Drosophila. In addition, the process has been documented and code provided so that it can be replicated in other systems. We load all of the raw data into an IGV viewer enabling the community to explore the data and compare the results for their own genes of interest (https://bio.rc.ufl.edu/pub/mcintyre/sex_specific_splicing/igv/index.html).

In a comparison between males and females, the classic Drosophila context dependent developmental paradigm, the fiveSpecies annotation combined with the empirical data identify context dependent alternative transcripts in adult head tissue in many more genes than previously suspected - including a large number of genes with both male and female-biased transcripts. This suggests that alternative transcript structure may be a common solution to sexual conflict in these species. High levels of conservation in sex bias in the *D. melanogaster* and *D. simulans* species pair support this inference, and the identification of conserved sex bias in transcript orthologs in *D. serrata* for genes outside the canonical sex determination pathway suggests that some sex bias is ancient while most sex bias is, as expected, rapidly evolving.

## Results

While differences in alternative splicing events can be inferred from short read or qRT-PCR based assessment of exons or splice junctions, short reads do not allow for the unambiguous resolution of isoforms^17^. Ambiguity in mapping due to sequence similarity between genes in gene families, and the divergence of paralogs between species, combined with uncertainty in the genome assemblies and annotations can all introduce bias^17, 18^. To mitigate these factors, one strategy for short reads is to rely on filtering strategies^19^. Long reads have much lower levels of ambiguity in mapping and allow splicing patterns for whole transcripts to be resolved^18^. In order to map each species’ data to their own reference genome, thereby limiting bias due to sequence divergence between the species, comparative evolutionary studies require the ability to identify the ‘same’ transcript between species. We develop a unified annotation of five species of Drosophila: *D. melanogaster, D. simulans, D. yakuba, D. santomea, and D. serrata* using a reciprocal liftover approach that leverages the individua; annotated transcriptomes of all five species (Figure 1A). While the current individual species annotations vary in the number of genes and transcripts, the fiveSpecies annotation for each species has a similar number of genes (∼18,000) and individual transcripts (∼77,000) on each set of genome co-ordinates (Supplementary Figure 1). Homologous transcripts are inferred across species by mapping the annotated transcripts from each species (source) to the other four species genome co-ordinates (targets). The map-based relationships are resolved using a network graph which identifies sets of homologous transcripts linked by mapping (Figure 1B). The unified genus level annotation has 15,996 genes with 56,370 homologous transcripts (Supplementary Figure 1). The annotation is validated with deep coverage long read data in males and females for all five species. The majority of transcripts are conserved in expression across the species. Replication in *D. melanogaster, D. simulans,* and *D. serrata* was used to identify quantitative differences in transcript usage between the sexes, and to evaluate the conservation of significant sex bias for transcripts in ∼10,000 genes in each of the three species.

**Figure 1.**
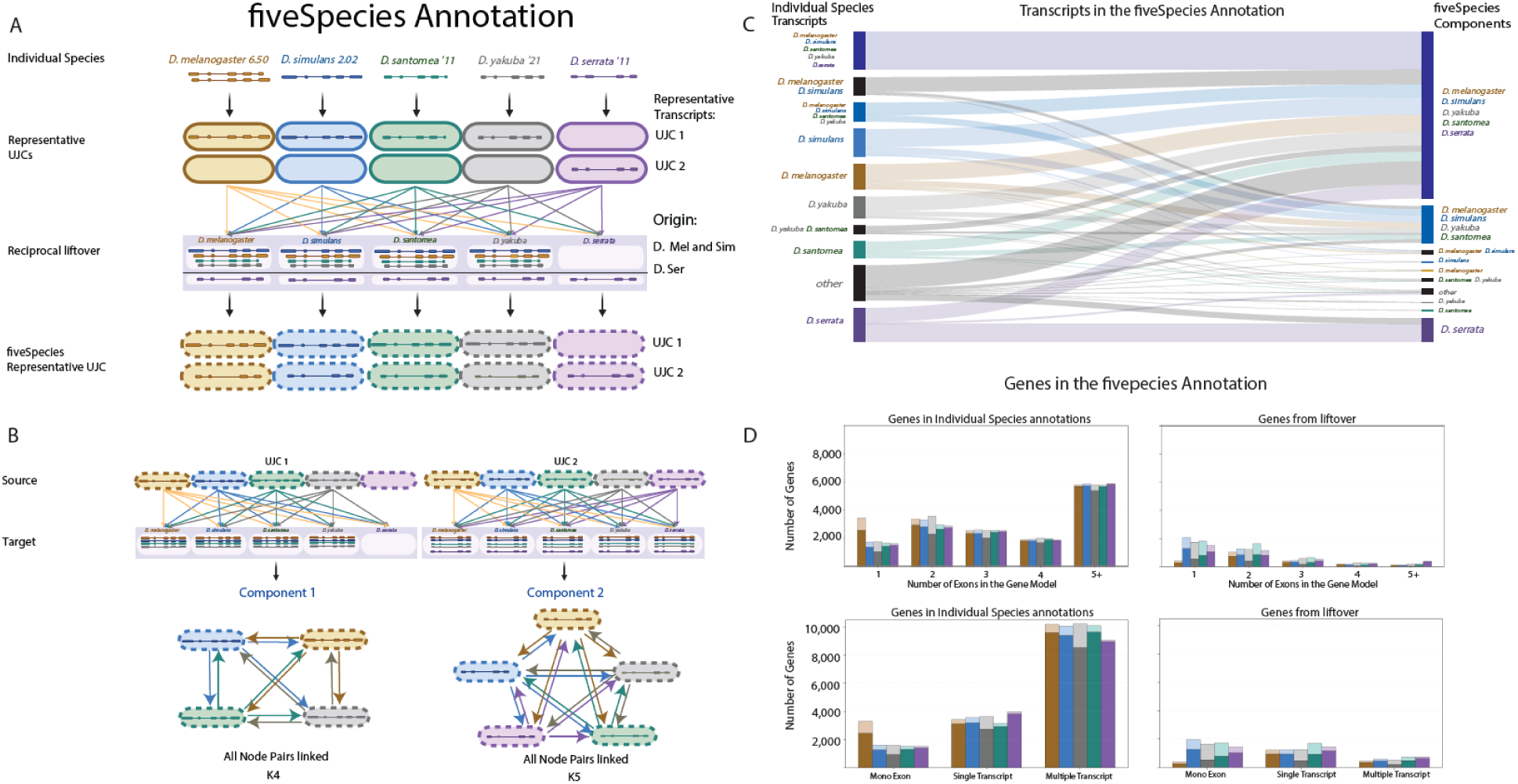
Unifying transcript annotations in Drosophila species. Panel A) Current individual species annotations for *D. melanogaster* (gold), *D. simulans* (blue), *D. yakuba* (gray), *D. santomea* (green) and *D. serrata* (purple) are mapped onto their respective genome coordinates. Transcripts that share the same internal intron structure (UJC) are annotated as a single representative transcript using the 5’ and 3’ most ends (*e.g. D. melanogaster*). A reciprocal liftover approach is used to map the individual species representative transcripts onto the other four species genomes. The resulting collection of mapped transcripts are summarized into representative transcripts (UJC) on each of the species genome coordinates to create the fiveSpecies annotations. Panel B) The fiveSpecies annotations are linked across coordinates using a network built from mapping representative transcripts from the individual species (source) to the other four (target) genomes and recording an edge when the mapping coincides with an annotated fiveSpecies transcript. Sets of linked transcripts are components in an associated network graph. Components represent likely orthologous transcripts, and the relationships can be described by the structure of the nodes and edges. When all nodes are linked reciprocally the node structure is K followed by the number of nodes. The structure of all 56,370 components (Supplementary Figure 1) can be described by their network. Panel C) The path from the individual species transcript annotations to the component identified in the fiveSpecies annotation (Supplementary Table: linking ujc_to_componentid). Ribbon color denotes the transcript source (left) and illustrates how individual species transcripts contribute to the shared fiveSpecies components (right). Panel D) The fiveSpecies gene annotations for *D. melanogaster* (gold, n=18,929), *D. simulans* (blue, n=18,981), *D. yakuba* (gray, n=18,885), *D. santomea* (green, n=18,891) and *D. serrata* (purple, n=18,195). Left, genes present in the individual species annotations. Right, genes lifted onto that species derived from another species’ annotation. The number of genes is the Y axis in all four bar plots. The X axis for the top row is the number of exons in the gene model, and the X axis on the bottom row separates genes annotated as mono-exon, single transcript or multi-transcript. The dark shading in all four bar plots indicates at least one of the fiveSpecies annotated transcripts is expressed in that gene.

### Identification of homologous/orthologous transcripts across multiple species

After reciprocal liftover, we summarize the set of transcripts with unique junctions on each genome and to identify homologous transcripts between species, we map each of the ∼77,000 fiveSpecies annotated transcripts from the source genome onto the four target genomes. When the source transcript maps to an annotated transcript in the target species an edge is recorded. By treating each source transcript on each set of co-ordinates as a node, we use the edges from the mapping function to build a network graph of components. Each sub-graph within the network is a component, and corresponds to a set of homologous transcripts (Figure 1B). Components are linked to each other in a second layer of the network analysis that uses gene identifiers. Of the 56,370 homologous transcripts, the majority (38,791, 69%) have homologs in all five species. This presents a large expansion of identified relationships of transcripts across species as only 8,796 homologs were present in the current individual species annotations for all species (Figure 1C, Supplementary Table 1, compare_origin_anno_xpress.csv). There are an additional 8,549 transcripts identified in the four species *D. melanogaster, D. simulans, D. yakuba,* and *D. santomea,* but not in *D. serrata* (Supplementary Figure 1).

### Annotation is enriched by the liftover in all species including D. melanogaster

Each species uniquely contributes an average of 5,700 transcripts to the fiveSpecies annotation (Figure 1D, Supplementary Table 2, transcript_model_counts.xlsx) and there are very few transcripts without identifiable homology to at least one other species (288, 219, 103, 114) in *D. melanogaster, D. simulans, D. yakuba,* and *D. santomea* respectively. Unsurprisingly, there are nearly three times as many transcripts added to *D. serrata* than in the current annotation, but there are also many more transcripts identified than currently annotated in *D. melanogaster*^5^. Transcripts added in the liftover are overwhelmingly in multi-exon/multi-transcript genes (*D. melanogaster-* gold bars, Figure 1D) and many of the ‘liftover’ transcripts are detected in the long read data (*D. melanogaster* - gold bars, right panel, dark shading, Figure 1D). Indeed, the fiveSpecies annotation describes the observed long read data substantively better than the current individual species annotations, for all five species. The proportion of reads that are associated with the annotation increases substantially for all samples in all species while the proportion of reads designated as intergenic or genic but unlinked to an annotated gene decreases (Supplementary Figure 3; Supplementary Table 3, sqanti_compare_native_vs_5species). There are ∼1,300 transcripts lifted onto *D. melanogaster* that are expressed in this study, demonstrating the value of the liftover even in this well studied model (Figure 2A; Supplementary Table 4, count_origin_anno_xpress_bySpecies.csv).

**Figure 2.**
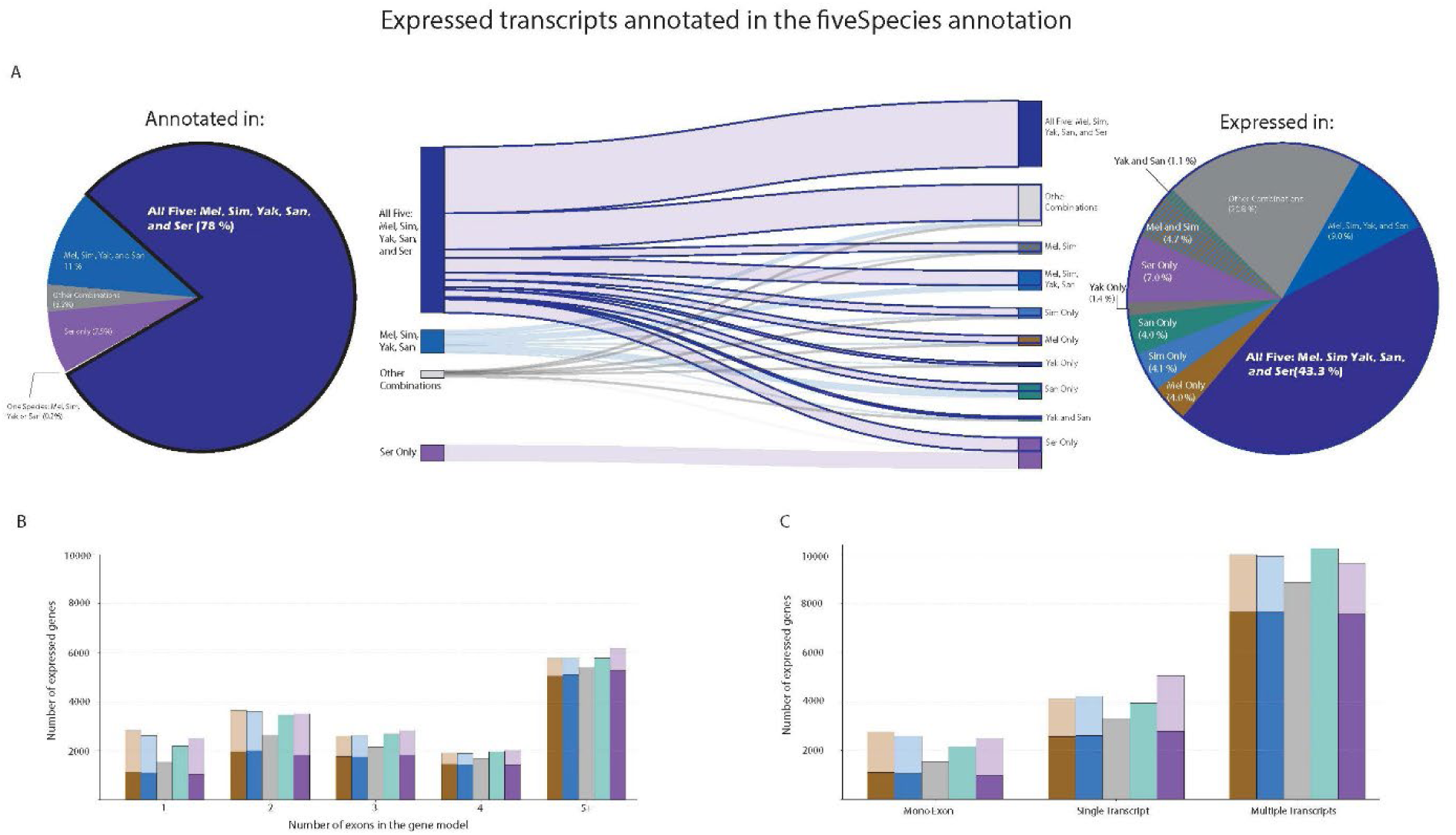
Empirical validation of the fiveSpecies annotation. Panel A) The component annotation (left pie chart) for expressed transcripts (right pie chart). Of the identified transcript orthologs, 38,791 are annotated in all five species (dark blue left pie chart). Outlined ribbons represent transcript orthologs expressed in all five species. Panel B) The number of expressed genes is on the Y axis and the number of exons in the gene model is on the X axis. *D. melanogaster* (gold, n=16,699), *D. simulans* (blue, n=16,501), *D. yakuba* (gray, n=13,412), *D. santomea* (green, n=16,061) and *D. serrata* (purple, n=16,993). Dark shading indicates genes with at least one fiveSpecies transcript analyzable for sex bias. *D. yakuba* and *D. santomea* have a single replicate and are not analyzed quantitatively for sex bias. Panel C) The number of expressed genes is on the Y axis and the X axis separates genes by annotated transcript model category. Mono-exon genes are defined as a single annotated exon, with no empirical evidence for additional exons. Single transcript genes have only one simple component. Colors and shading as in Panel B.

### Most gains in the liftover are from adding transcripts to genes already annotated in the species

The value of the fiveSpecies annotation for investigating the conservation and evolution of alternative splicing is largely in adding transcripts to existing annotated genes, as illustrated by the well-known alternatively spliced *Drosophila* gene, *Sxl* (*Sex lethal*). *Sxl* has only one annotated transcript model in the *D.serrata, D. yakuba* and *D. santomea* annotations, although molecular evidence for at least two transcript models, female and male (with and without a male specific exon) are expected in each of these species with biological implications for sex determination ^38, 39^. The fiveSpecies annotations for *Sxl* contain transcripts both with and without the male-specific exon in all five species as well as transcripts containing exon Z, which is expressed in both sexes (Supplementary Figure 4, ^40^). The improvement in the *D. melanogaster* annotation is illustrated by component 14,438, of the gene *NK7.1* (Supplementary Figure 2). Individual species annotations in *D. serrata* and *D. simulans* were the basis of the fiveSpecies annotation of this transcript and it is expressed in *D. melanogaster* (Supplementary Figure 2).

### The network approach facilitates transcript ortholog identification

Approaches identifying orthologous exons are frequently based on multiple alignment and phylogenetic analyses^21^. Generally, this is coupled with manual curation to resolve ambiguity in junctions and splits/fusions between exons^41^. In contrast, by linking transcripts using mapping functions to create edges between species, and using a network formed from these edges, no upfront resolution to the exon structure between species is needed. Instead, the relative structure of the transcript can be annotated with a digital binary and differences in structure summarized and used to identify ambiguity in the structure. The associated empirical data can then be used to resolve ambiguity. Individual species data are mapped to their respective genomes and compared to the annotation to resolve exon splits/fusions. For example, in *Sxl*, *D. yakuba* and *D. santomea* have more exons (16) than *D. melanogaster* and *D. simulans* (15) (Supplementary Figure 4A). An all-by-all blastn of exons verifies that one exon in *D. melanogaster* (X: 7074549-7079367) matches two separate regions in *D. yakuba* (NC_052526.2: 14852264-14856600 and NC_052526.2: 14851526-14852189) with a 75 bp gap between the two *D. yakuba* regions. Alignment of the underlying genomic DNA in a blast2seq reveals a high level of sequence divergence between the species with a 15 bp indel, followed by 24 bp with 22 matches and 2 mismatches, and an additional 35 bp with poor sequence similarity across species. Our computational pipeline annotates each empirically observed long read for introns and additional exons relative to the gene model (github.com/McIntyreLab/TranD/tree/main/utilities/id_erp.py). In this case, the observed long read transcripts contain an ‘intron retention’ for both species, supporting transcription across the putative ‘split-exon’ region in *D. yakuba* and *D. santomea.* These data indicate a single transcribed region with a region of sequence divergence between the species (Supplementary Figure 4B, see zoom out inset). Our annotation pipeline enables the further development of computational solutions to resolve putative splits/fusions at scale, where previous efforts, by necessity, were almost completely manual. In the same way we can resolve ambiguity in splits/fusions, we can resolve ambiguity in splice junction identity when the liftover from different source genomes produces similar but not identical splice junctions on the target genome. For example, in component 14,438, there are multiple transcripts in the fiveSpecies annotation for the *D. serrata* genome and a single transcript in the other four species. All the *D. serrata* transcripts have the same pattern of exon inclusion but vary in the splice junctions. The empirical data from *D. serrata* corresponds to one of the annotated transcripts in the set. We consider the transcript supported by the data in *D. serrata* to be the likely ortholog and the resulting annotation now has a single node per species. Applying this logic to the components annotated in all five species, but with multiple nodes in one or more species we identified the components with one expressed node per species and considered these together with the components with a single annotated node in all five species, as orthologs and there were 23,352 of these (Supplementary Figure 1).

### There are potential orthologous novel transcripts

Long read experiments are acutely sensitive, identifying many potential novel transcripts^28, 42^. We define novel transcripts as those that are not in the annotation and for which there is putatively a distinct biological function from the set of annotated transcripts for the gene (a novel exon pattern). We further limit our consideration of novel transcripts to those in well-expressed (100 reads or more) genes lacking evidence in the data for annotated transcripts (cases where the fiveSpecies annotation explains less than 1% of the observed data). We consider transcripts with >20% of total reads in the gene as putatively novel transcripts^43^. There are 263 transcripts that meet this strict definition across all five species. Some of the potentially novel transcripts were identified in multiple species among orthologous loci (Supplementary Table 5, putative_novel_transcript_link_orthologs.csv). For example, a novel transcript in the *D. melanogaster* genes FBgn0034928 (CG13562 - predicted to enable triacylglycerol lipase activity) and FBgn0037679 (*Aduk*) have observed transcripts in the long read data for *D. yakuba*, *D. santomea* and *D.serrata.* The list of these transcripts is provided (Supplementary Table 6, count_putative_novel_transcripts.csv; Supplementary Figure 5)

### Expressed transcripts are conserved with potential lineage specific transcript structure in the outgroup

We detected 49% of the annotated transcripts as expressed in head tissues (27,876/56,370). The vast majority are expressed in all five species (78%) (Outlined ribbons, Figure 2A) or in four of the five species (11% - *D. melanogaster, D. simulans, D. yakuba, D. santomea).* Among the expressed transcripts, 43% (8,704) are expressed in all five species, and there are relatively few expressed only in the *D. melanogaster* and *D. simulans* branch (1,385) and even fewer expressed only in the *D. yakuba, D. santomea* branch (515) with 30% expressed in various combinations of species making it difficult to rule out that these transcripts are present but not observed due to sampling variance^44^ (Figure 2A - right pie chart; Supplementary Table 4, count_origin_anno_xpress_bySpecies). There are 4,062 transcripts expressed only in *D. serrata* (7.2%), and these are potential candidates for divergence in *D. serrata*. This combined evidence leads us to estimate that at least 73% and perhaps up to 90% of expressed transcripts are conserved. We cannot rule out the possibility of expressed orthologous transcripts for the transcripts detected only in *D. serrata,* but sequence divergence may have impeded the liftover approach.

### Transcript diversity is suggestive of functional diversity

For genes with only a single exon, transcript variation is limited to potential 5’ and 3’ UTR length and poly-A variation. This type of transcript diversity is associated with differences in regulation, such as localization and transcript stability; as only one protein product is made with a single combination of functional domains. mRNA seq does not capture UTRs accurately and therefore we do not separate transcripts based on the 5’ and 3’ ends. The fiveSpecies annotation contains approximately 3,000 mono-exon genes (∼15%). For genes with multiple exons, there is the potential for different transcripts that vary in exon length and combinations, and a greater possibility for evolutionary changes to modulate expression and/or products in different contexts with some degree of independence – in other words, there is relaxed constraint and enhanced capacitance for transcript evolution. The genes and transcripts lifted across species are overwhelmingly multi-exonic in their structure (Supplementary Figure 3) and the fiveSpecies annotation suggests 50% of genes have multiple transcripts (Figure 1D), an increase over the current annotation of 40%. We categorized the annotated transcripts of multi-transcript genes to assess structural differences based on alternative donor/acceptor sites, intron retention, and alternative first, last or internal exons; while alternative donor/acceptor sites and first exons were the most prevalent categories, there are substantial levels of intron retention and alternative last or internal exons (Figure 3C-D). Each of these can be associated with functional differences among transcripts^45, 46^.

**Figure 3.**
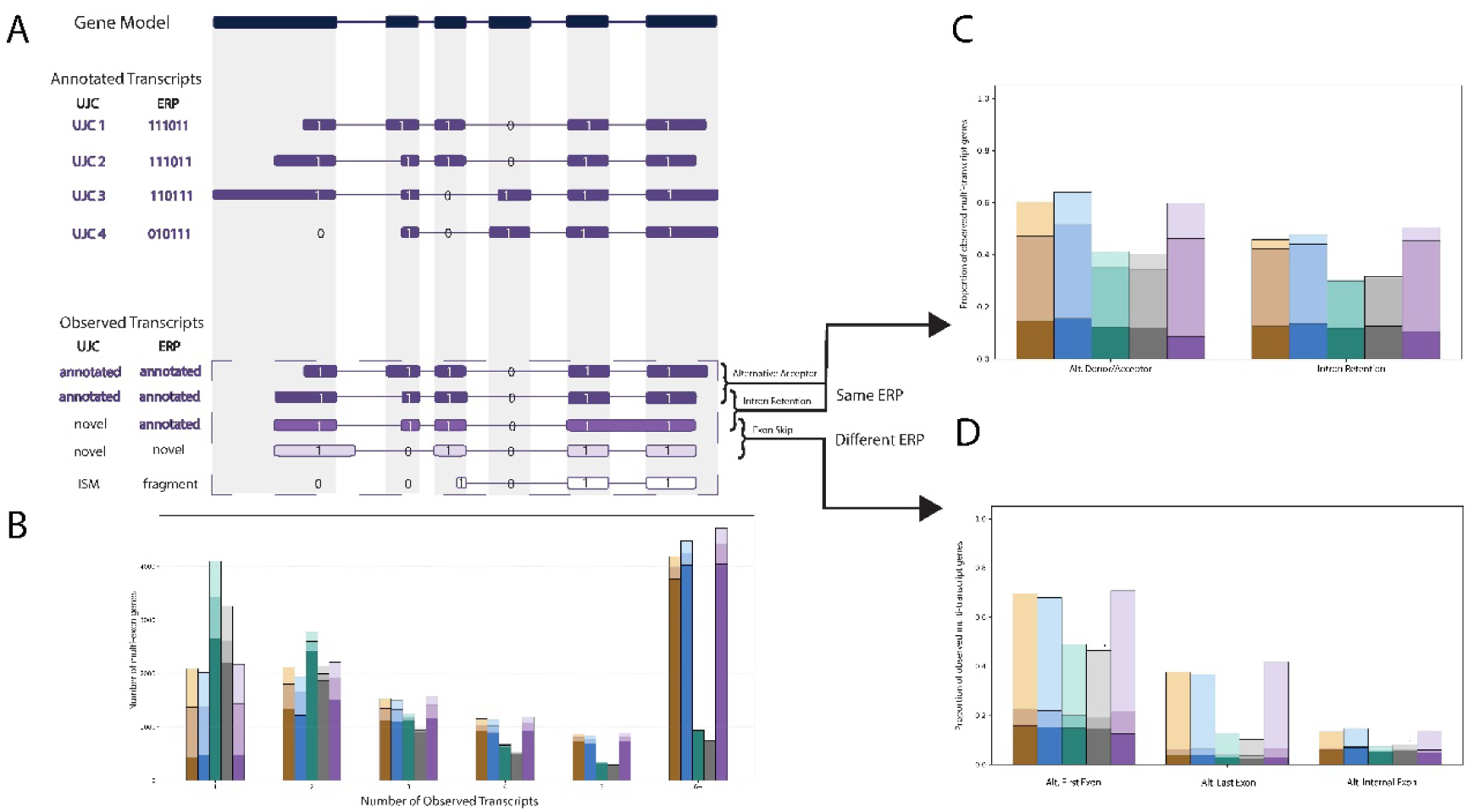
Alternative splicing in Drosophila. Panel A) Exon patterns are determined by binary indicators that identify the exons present in each transcript relative to the gene model. Two transcripts that share the same exon pattern but differ in their junctions have either alternative donor/acceptor variation and/or an intron retention. Alternative first (5’) and last (3’) exon inclusion, and exon skipping are determined by comparing patterns between transcripts. These three categories are not mutually exclusive. Observed transcripts may be consistent with annotated transcripts, consistent with annotated exon patterns, or novel. Transcripts that are a subset of annotated exons (incomplete splice match, ISM) are considered fragments. Panel B) The Y axis is the number of observed multi-exon genes in *D. melanogaster* (gold, n=14,626), *D. simulans* (blue, n=14,380), *D. yakuba* (gray, n=12,338), *D. santomea* (green, n=14,740) and *D. serrata* (purple, n=15,648). The X axis is the number of observed transcripts. Transcript fragments are not included. Dark shading indicates at least one observed transcript is annotated. Medium shading indicates that, of the genes without an observed annotated transcript, there is at least one transcript with an annotated exon pattern. Light shading indicates all observed transcripts are novel. Panels C+D) The Y axis is the proportion of genes with multiple observed transcripts with at least 10 reads: *D. melanogaster* (gold, n=9,868), *D. simulans* (blue, n=9,887), *D. yakuba* (gray, n=4,685), *D. santomea* (green, n=5,995) and *D. serrata* (purple, n=10,572) with at least one annotated transcript (dark shading), or at least one transcript with an annotated exon pattern (medium shading), or at least one full length novel transcript (light shading). Panel C) The number of observed multi-transcript genes with at least two transcripts that share an exon pattern: *D. melanogaster (*gold, n=6,433), *D. simulans* (blue, n=6,801)*, D. yakuba* (gray, n=2,821)*, D. santomea* (green, n=3,538)*, D. serrata* (purple, n=7,278). The X axis separates alternative donor/acceptors and intron retentions. These categories are not mutually exclusive. Panel D) The number of observed multi-transcript genes with at least two transcripts with different exon patterns: *D. melanogaster* (gold, n=5,508), *D. simulans* (blue, n=11,777)*, D. yakuba* (gray, n=9,146)*, D. santomea* (green, n=10,940)*, D. serrata* (purple, n=12,945). The X axis separates alternative first (5’) and last (3’) exons and exon skips. These categories are not mutually exclusive.

### Most expressed genes have multiple expressed transcripts

The brain is a tissue made up of many cell types with the expectation that there should be diversity of isoforms among these cell types^47–49^. With ∼200 million mapped long reads per species, we were able to reliably evaluate more than 10,000 genes for multiple transcripts in *D. melanogaster*, *D. simulans* and *D. serrata.* With ∼100 million reads in *D. yakuba and D. santomea,* we were able to evaluate 6-8 thousand genes for multiple transcripts (Figure 2B, 2C). Many studies using long read RNA-seq report identifying large numbers of novel transcripts^50, 51^. Short read RNA-seq studies often show 3’ bias^52^ and long reads are no exception^53^. However, unlike with short reads, likely transcript fragments in studies with long reads can be identified, as long as there is a reasonably complete annotation^28, 29^. Using the fiveSpecies annotation as the reference, reads were classified in the following mutually exclusive categories: originating from a likely transcript fragment (Incomplete Splice Match, ISM), consistent with an annotated transcript (Full Splice Match, FSM), having the same exon structure as an annotated transcript (sharing an Exon Region Pattern, ERP), or novel (Figure 3A).

### Expressed alternative transcripts indicate a level of complexity in the Drosophila transcriptome previously underappreciated

Excluding likely transcript fragments, and lowly expressed transcripts (fewer than 10 reads), and the transcripts with an annotated ERP, we observed many novel expressed transcripts. Indeed, once we include novel transcripts in evaluating genes for the expression of multiple transcripts, 75% or more of genes can be considered to have multiple expressed transcripts (Figure 3B). Using a faster implementation of the distance metrics presented in TranD^23^, we compared all expressed full-length transcripts in the same gene pairwise. For pairs that share an exon pattern, differences between transcripts in junctions and intron retentions are identified. For pairs that differ in their exon structure, we use a set operation on the ordered set of exons to identify alternative first and last exons as well as exon skips. We evaluate expressed alternative transcripts at two levels of stringency-the first level of moderate stringency includes all transcripts expressed that are not likely fragments (Supplementary Figure 3), and at the second level of high stringency we restrict comparisons to only transcripts with at least 10 reads (Figure 3C-D). While novel alternative first exons may be a technological artifact, alternative skips and alternative last exons are likely true variants. The existence of high rates of exon skips at both levels of stringency reveals a high level of complexity in the *Drosophila* transcriptome.

### Sex bias is pervasive and rapidly evolving

We identify sex bias in transcript usage in nearly 40% of genes (Figure 4C). There is conservation of bias in genes primarily in the *D. melanogaster*, *D. simulans* species pair but also extending to *D. serrata* (Figure 4C). At the gene level, conserved bias may be explained by homologous or orthologous transcripts, in which case sex-differences in transcript usage are conserved. However, there may be similar selective pressure driving the evolution of sex bias in the same direction in multiple species, and the response may not always produce bias in the homologous transcripts. As a result, there may be different transcripts biased in the same direction for the same gene. We identified 605 transcript orthologs conserved for sex bias in the *D. melanogaster-D. simulans* species pair, a far higher number than expected based on the literature-but far short of the marginal bias rates observed at the gene level, in each species. However, even in the closely related *D. melanogaster* and *D. simulans* species pair there is evidence for shifts in sex bias for individual transcripts (Figure 4D), suggesting differences in selective pressures or that that similar selective pressures may result in different solutions at the individual species level.

**Figure 4.**
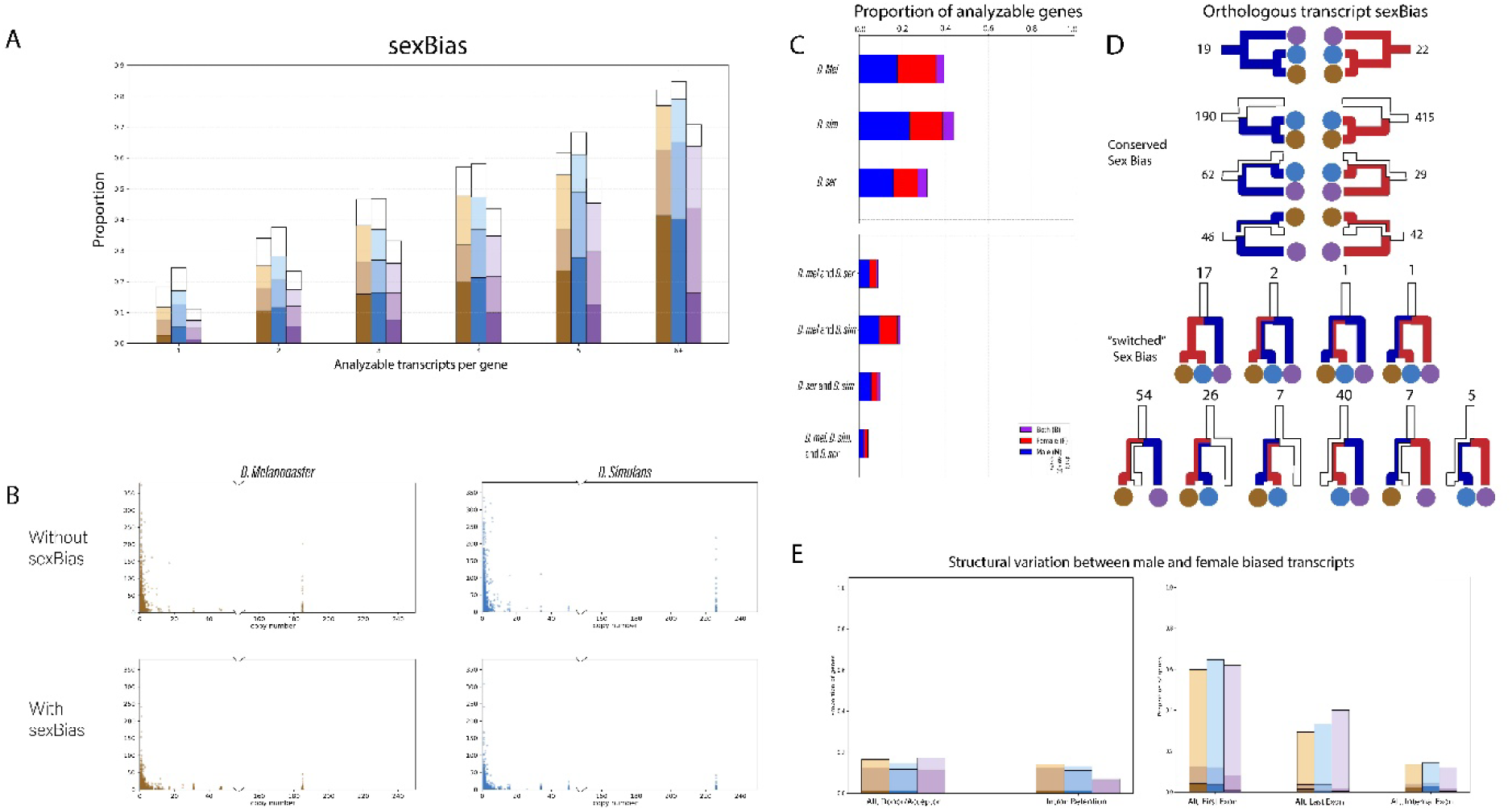
Context dependent alternative splicing: sex bias in Drosophila. Panel A) For all genes evaluated for sex bias in *D. melanogaster* (gold, n=11,313), *D. simulans* (blue, n=11,371), and *D. serrata* (purple, n=11,360) the proportion with at least one annotated sex biased transcript (dark), or at least one annotated exon pattern (medium), or at least one novel exon pattern (light) or a transcript fragment (white box). Multi-transcript genes are more likely to exhibit sex bias (p<<0.0001). Panel B) The Y axis is the estimated number of transcripts per gene. The X axis is the number of genes in the gene family (Demuth and Hahn 2009). The top row is genes with sex bias and the bottom row is genes without statistical evidence for sex bias. Left, *D. melanogaster* and right, *D. simulans*. All transcript fragments are excluded, and the number of distinct exon patterns in the remaining set of transcripts is used as an estimate for the total number of distinct transcripts. In all cases there is a negative relationship between the number of transcripts and number of duplicates. *D. yakuba* also has a negative relationship between number of transcripts and number of duplicates (Supplementary Figure 6). Panel C) The proportion of analyzable genes with sex bias in male transcripts (blue), female transcripts (red) or both (purple) in each of the three species, and with conserved patterns of bias between the species pairs and for all three species (Supplementary files: gene_summary_<SPECIES>.csv, merged_fivespecies_genesummmary.csv) Panel D) Sex bias among orthologous transcripts. Red signifies female bias and blue signifies male bias. (Supplementary file: Component_summary.csv). Panel E) Alternative splicing patterns for multi-transcript genes with both a male and female biased transcript: *D. melanogaster* (gold, n=410), *D. simulans* (blue, n=608), and *D. serrata* (purple, n=493). Dark shading indicates the variation in sex bias is between annotated transcripts, medium shading indicates the variation in sex bias is between transcripts with an annotated exon pattern and light shading indicates the variation in sex bias is between novel transcripts. Left, male and female biased transcripts share an exon pattern: *D. melanogaster* (gold, n=92), *D. simulans* (blue, n=121), and *D. serrata* (purple, n=93). The X axis separates alternative donor/acceptors and intron retentions. These categories are not mutually exclusive. Right, male and female transcripts have different exon patterns: *D. melanogaster* (gold, n=208), *D. simulans* (blue, n=339), and *D. serrata* (purple, n=285) The X axis separates alternative first (5’) and last (3’) exons, and exon skips. These categories are not mutually exclusive.

### Sex bias in head tissue is associated with the developmental sex-determination pathway

While the key regulators in the sex determination pathway compose a splicing cascade, they can also have independent targets with additional interactions among genes in the pathway^4, 54–67^ (Figure 5A). For example, different neurons in the brain individually express Doublesex (Dsx) or Fruitless (Fru) or they can be co-expressed^68–70^. There is evidence for sex-specific splicing in splicing factors and, together with independent branches of the pathway, this suggests the pathway regulates sex-specific isoforms of additional targets^13, 71, 72^. Genes with sexually dimorphic transcripts are likely candidates for sexually dimorphic phenotypes and may be regulated by the sex determination pathway. For example, the gene *Glutamine synthetase 2* (*Gs2)* was previously identified as a direct target of Sxl^73^, our data show conserved male-biased transcripts for this gene. In total, we identify 22 male and 19 female biased transcripts where sex bias is conserved with the outgroup *D. serrata,* including transcripts of Glutamine synthetase 2 (*Gs2)* and *ciboulot (cib)*. *cib* encodes a homolog of an actin binding protein in the thymosin beta gene family, with a role in brain development and we identify a transcript with conserved male bias in all three species (Figure 5B; ^74^). These genes have not been implicated in alternative transcription or sex bias in the head, although previous studies showed *Gs2* is male-biased in whole adults and it is associated with glutamatergic signaling and brain development – similar to *cib*^73, 75^. To understand the correspondence of sex-biased transcription to likely targets of the sex determination pathway more broadly, we examined a suite of prior studies. In our previous study we identify candidates for incorporation in the sex determination pathway in *D. melanogaster*, and of those we can evaluate in both studies, 39 have sexually dimorphic transcripts in this study^76^. Genes regulated by or downstream of *tra* are also enriched among genes with sex-biased transcripts in our study, with as much as double the proportion expected by chance (Supplementary Table 7, list_compare_percent2.csv ^77, 78^), as are those downstream of *dsx* and in males downstream of *fru* (Supplementary Table 7, list_compare_percent2.csv ^60, 79^).

**Figure 5.**
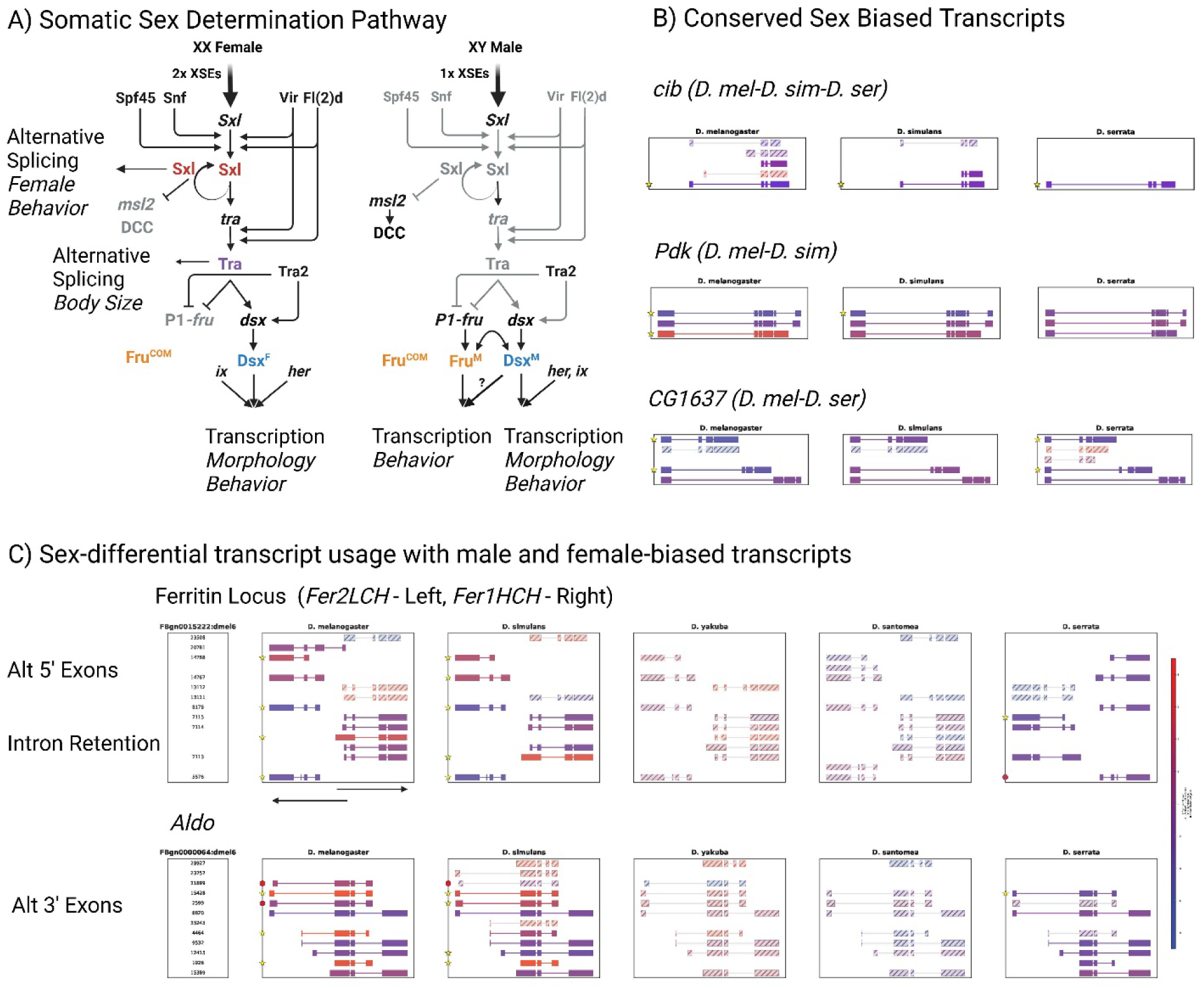
Sex differences in alternative transcripts. Panel A) The regulation of alternative transcripts by the somatic sex determination pathway in *D. melanogaster*. There are four central regulators with one or more alternative transcripts, and distinct cofactors, that form a splicing cascade. Sex lethal (Sxl) is an RNA binding protein that regulates alternative splicing, translation and polyadenylation switching. Transformer (Tra) is part of a splicing enhancer complex that regulates alternative splicing of the genes encoding the Doublesex (Dsx) and Fruitless (Fru) transcription factors. Each regulator also defines an independently acting branch of the pathway. Sxl can act independently of its target Tra, in regulating dosage compensation and female behavior. Tra can act independently of Dsx and Fru. There are independent targets of the transcription factors Dsx and Fru, and evidence for contexts where they interact. Other interactions have been suggested. Panel B) Conserved patterns of sex-differential transcript usage. Gene models are shown to scale for the corresponding genomic reference from 5’ to 3’. A transcript of the gene *ciboulot (cib),* corresponding to *cib*-RA/RC in *D. melanogaster,* shows conserved male bias between *D. melanogaster*, *D. simulans,* and *D. serrata*. A transcript of the gene *Pyruvate dehydrogenase kinase (Pdk),* Pdk-RB, shows conserved male bias only between *D. melanogaster* and *D. simulans*. Transcripts of the gene *CG1637* show conserved male bias (CG1637-RA/RE and CG1637-RB/RD) between *D. melanogaster* and *D. serrata*, but not *D. simulans.* Transcripts with yellow stars indicate significant sex bias at a nominal p < 0.05. The set of transcripts sharing the same exon pattern were also assessed for sex bias with filled circles indicating a significant t-test at a nominal p < 0.05 level. The fill color of transcripts and of circles show the degree of sex bias as indicated by the effect size estimated by the t value. Data are in supplementary files datafile_jxnHash_<SPECIES> w_annoFlag_ERP_ESP_info_strCat_flagNovel_02amm and can be viewed in an IGV browser at https://github.com/McIntyre-Lab/igv_4_sex_specific_splicing. Panel C) Gene models are shown to scale for the corresponding genomic reference from 5’ to 3’, with the exception of *Fer2LCH*. Transcript annotations indicating sex bias are as described for Panel B. Top, male and female alternative transcripts expressed from the ferritin locus in *D. melanogaster* and *D. simulans*. All data shown in Supplementary Figure 7. Alternative transcripts of *Fer1HCH* differ in the presence of an intron retention between exons 1 and 2 that contains an Iron Response Element (IRE), which in this case is expected to inhibit translation and ultimately impact iron level homeostasis. *Fer2LCH* has both male and female-biased transcripts, with Fer2LCH-RD and Fer2LCH-RE showing conserved male bias in *D. melanogaster* and *D. simulans*. In *D. serrata*, most transcripts show common expression with similar expression levels in males and females, but both genes do have at least one male-biased transcript. Bottom, multiple female biased transcripts of the gene *Aldolase* (*Aldo*) are conserved between *D. melanogaster* and *D. simulans*. Commonly expressed transcripts, purple, that are expressed in both males and females are typically conserved in their expression pattern between all three species, but female biased transcripts are not observed in *D. serrata*.

### The list of tissues showing sex differences in transcript structure is expanded by these data

For example, *Aldolase* (*Aldo*) is expressed in heads at a similar level in males and females (flybase – FlyAtlas2^80^). Evidence for sex differences at the transcript level, which may vary across genotypes and tissues, has been noted previously in *D. melanogaster*, where there may be variation across tissues and genotypes^13, 81^. The alternative transcripts encode conserved (human-fly) alternative C-terminal regions that are known to impact enzymatic function of the protein products^82–84^. Alternative transcripts with alternative C-terminal regions show conserved differences in male/female usage between *D. melanogaster* and *D. simulans* (Figure 5C). A similar pattern of alternative C-terminal regions and conservation of sex differences in transcript usage was observed for the gene *Pdk*, *pyruvate dehydrogenase kinase* in *D. melanogaster*: Pdk-RB is male-biased, Pdk-RA is female biased, and Pdk-RD is common. Both *Pdk* and *Aldo* have been studied in the context of sex-differences in midgut expressed genes regulated by gonad to midgut JAK-STAT signaling and impact feeding behavior in males^85^.

### We observe novel sex differences in transcript usage

For example, in the genes encoding the two subunits of Drosophila ferritin, *Ferritin 1 heavy chain homologue* (*Fer1HCH*) and *Ferritin 2 light chain homologue* (*Fer2LCH*). Ferritin has a conserved role in iron-binding and storage, and both genes express multiple alternative transcripts and they partially overlap one another, so may share regulatory regions^86–89^. At the gene level these genes show male bias in *D. melanogaster* and *D. simulans*^80, 90^. We find that in both *D. melanogaster* and *D. simulans* the transcript of Fer1HCH-RA, is strongly female biased, whereas the other transcripts are similarly expressed (Figure 5C). The transcript Fer1HCH-RA retains an intron which contains an Iron Response Element (IRE; ^87^). In fact, we find sex differences in transcript usage for both genes at this locus that are not observed in *D. serrata*. In *Drosophila* there are known sex differences in iron homeostasis^91^. Interestingly, this gene has also been identified as potentially evolving due to sexual conflict in stalk-eyed flies, with male bias evolving in duplicated copies^92^.

### There is striking evidence for alternative male and female biased transcripts expressed from the same gene

One of the predictions for the resolution of intralocus sexual conflict is the evolution of alternative male and female transcripts expressed from the same gene. The textbook example of alternative male and female transcripts is somatic sex determination in Drosophila. The sex-determination gene regulatory network (SD GRN) consists of a hierarchical splicing cascade^4, 69, 93–100^. Expression of sex specific transcripts of *Sxl, fru* and *dsx* is conserved in Drosophila^71–79^. Alternative splicing of *Sxl*, *transformer* (*tra*) and *transformer 2* (*tra2*), lead to differences in protein isoforms at several downstream loci, including Dsx and Fru, resulting in somatic sexual differentiation during development^54–62, 67, 94^ (Figure 5A). We observe the expected transcripts for the male specific, female specific, and commonly expressed transcripts for Sxl^40, 101^ (Supplementary Figure 5) and there are a number of genes with sex-dimorphism in transcript usage suggested in the literature that are conserved between *D. melanogaster* and *D. simulans* (Figure 5; e.g. *Aldo*, *Pdk*, *Desat1* – See Supplementary Figure 8)^13, 81, 102, 103^. There is also an enrichment in this study for genes previously identified as having likely sex bias in alternative exons^13, 71, 72^. There are 410 genes in *D. melanogaster*, 608 genes in *D. simulans* and 493 genes in *D. serrata* with evidence of both male and female biased transcripts, suggesting that alternative transcripts may contribute more widely to the evolution of conflict resolution than previously appreciated. These include genes where functional differences between transcripts are established, such as *Aldo*, where both the male and female biased transcripts are conserved in the *D. melanogaster*, *D. simulans* species pair. There are 118 orthologous genes with both a significantly male biased and a significantly female biased transcript and some additional genes where bias trends are similar between the species.

### There are large differences in transcript structure between male and female biased transcripts of the same gene

By comparing exon patterns between the male and female biased transcripts we identify the types of structural differences between them (Figure 4E, Figure 5C, Supplementary Figures 2, 4, 7, 8). There are about twice as many of these genes with different exon usage patterns compared to transcripts with shared exon usage, with many fewer alternative donor/acceptor sites compared to intron retention. For genes with alternative exon usage patterns in the male and female biased transcript, alternative first or last exons are prevalent with some evidence for exons skips (Figure 4), similar to differences between transcripts overall (Figure 3). In addition, genes with sex-biased transcripts are enriched for genes with predictions of alternative exon cassette usage (Supplementary Table 7 list_compare_percent2.csv; ^104^). The structural differences between the male and female biased transcripts are consistent with functional differences, although the association of specific alternative isoforms with functional differences that resolve conflict is a multi-factorial process involving evolutionary history, functional impacts and fitness effects^105, 106^.

### Gene duplication and alternative splicing are inversely related

Duplication and alternative splicing can lead to functional divergence, and an inverse relationship has been found between gene family size and the number of alternatively spliced isoforms^22^. This trend has been documented across species such as humans, mice, flies, zebrafish, and frogs^6, 107^. We find a negative relationship between the number of transcripts and the number of duplicate genes in *D. melanogaster, D. simulans* and *D. yakuba,* the three species in this study with estimation of gene duplicates^108^ (Supplementary Figure 6).

### Gene duplication status is not associated with sex bias

We also examine the correspondence of transcript number and gene duplication to sex bias in *D. melanogaster* and *D. simulans* (Figure 4B), where we were able to statistically test for quantitative differences in sex bias for individual transcripts. Significant sex bias is said to be present for the gene if at least one transcript is statistically significant for sex bias. We compare the relationship between transcript number and duplication in genes with evidence for sex bias (top row) and those without evidence for sex bias (bottom row) (Figure 4B), and note that the overall trend is unaffected by sex bias. There is a cluster of 185 genes in *D. melanogaster*, 226 in *D. simulans*, and 329 in *D. yakuba* and some of these genes also have multiple transcripts (Figure 4B). In *D. melanogaster* and *D. simulans* there is evidence for sex bias among the genes in this cluster. The cluster contains many ribosomal genes. There is no association between sex bias (yes/no) and the status of a gene as duplicated (yes/no) (P = 0.41). Examining differences between mono-exon genes, single transcript genes and multi-transcript genes, there is a significant difference in the frequency of sex bias across these categories (P < 0.0001). There are 52 mono-exon or single transcript genes in *D. melanogaster* (no multiple transcripts present) for which we have information on the duplication status. Two of these are sex-biased genes, both of which have at least one duplicate.

### The primary mechanism for the resolution of sexual conflict in these data is most likely to be alternative splicing

In contrast to duplicated genes, sex bias is more common in genes with many (6+) transcripts and depauperate in genes with only 1 transcript (Figure 4A). As a caveat, since we test each transcript and we also test groups of transcripts with the same exon pattern, genes with more transcripts will have more tests conducted increasing the likelihood of observing spurious sex bias. The proportion of genes with sex bias is similar for genes with 2 - 5 transcripts, and genes with one transcript have limited evidence for sex bias, suggesting the observation is not entirely a statistical artifact. Overall, the predominant evidence suggests that while the resolution of sexual conflict can involve sex-specific or sex-biased expression of duplicates, duplication is not the sole mechanism. Alternative splicing may be the more common evolutionary solution to sexual conflict in the tissues evaluated in this study.

## Discussion

### The fiveSpecies annotation enables comparative evolutionary studies of alternative transcript structure in Drosophila

The Drosophila fiveSpecies annotation provides a framework for comparative evolutionary studies of alternative transcript structures in the genus Drosophila, with 15,996 genes and 56,370 transcripts, of which 23,352 are likely orthologous transcripts. The identification of multiple homologous transcripts per gene supports comparative studies of conservation and evolution of alternative transcript structure in multi-transcript genes – despite differences in genome quality^3, 6^. Short reads can identify important splicing events, but long reads are needed to resolve transcript structure^18^. Overall transcript structure is likely to be important for functional differences, as comparisons of observed transcript structures in long read data from eukaryotes demonstrates that alternative exon usage is likely to co-occur with other structural changes in the transcripts^72^. Comparative studies of short reads often rely on cross-species mapping, a process that is inherently biased although new approaches to filtering may ameliorate some of those effects^19^. In this quantitative long read RNAseq study of male and female head tissue in five species in the Drosophila genus, data are mapped to their own species and transcript homology is used to compare expression patterns across species thus avoiding several layers of known bias and the need to identify homologous exons and specify a principle isoform. Using the fiveSpecies annotation and ∼600 million long reads, we estimate that 73%-90% of orthologous transcripts are conserved in expression and infer conservation of expression for multiple transcripts in 75% of the genes studied. Despite this dramatic improvement in annotation, some genes are still poorly annotated. The long reads reveal a transcriptional complexity far greater than previously imagined, with 75% of genes expressing multiple structurally distinct alternative transcripts in all five species. Our binary scoring of the transcript structure enables us to rapidly identify transcripts that share the same structure as existing annotated transcripts, as well as identify structural differences between any pair of transcripts regardless of annotation. This summary can be used to capture patterns in differences of transcript structure.

### Transcript complexity in Drosophila has been drastically underestimated

The fiveSpecies annotation provides substantial improvement to all of the individual species’ annotations, and transforms the annotation of *D. serrata*, an emerging but less well studied model, for sexual selection, conflict and aging^34, 35, 109–112^. We are able to rapidly identify the known set of transcript orthologs for the sex determination genes *Sxl*, *tra* and *dsx* in *D. serrata* that are absent in the current annotation. The fiveSpecies annotation also improves the annotation of *D. melanogaster*, with support in the data for more than 1,000 transcripts lifted from other species onto the *D. melanogaster* genome. Annotated transcripts are typically heavily curated and are shaped by the research focus of the community contributing to genomic and transcriptomic studies^113^. Transcripts with evidence from high-throughput data are not always included in annotations, particularly when there are many variants and both the structure of the transcript and their functional relevance is unclear. Our data-driven comparative evolutionary approach provides evidence for the conservation of alternative transcripts and supports further study, arguing for inclusion of conserved structural variants in the annotation even if there is no immediate understanding of the functional consequences of these variants.

### Alternative splicing is a likely mechanism for the resolution of sexual conflict in head tissues

Partial duplications or other mutations that impact splicing patterns can lead to novel transcript isoforms^114, 115^. Once multiple isoforms arise, their products are expected to become differentiated via evolutionary processes, leading to distinct functions and roles relative to the ancestral state^6, 12, 116^. Context is a fundamental aspect of this process, contributing to the distinction between gene copies or transcript isoforms and the repertoire of possible gene interactions based on patterns of co-expression^6, 9^. Differentiation of gene duplicates or alternatively spliced isoforms across lines of sex, tissue and cell-type can evolve when fitness optima differ between sexes leading to conflict^105^. The relationship between conflict and sex differences in gene regulation results from the resolution of existing sexual conflict by the evolution of sex differences. For example, imagine that a single transcript is expressed from a gene and the most frequent allele benefits male fitness but reduces female fitness. This might be resolved in a number of ways, but one possibility is the evolution of sex differences in transcript usage, allowing decoupling of expression; each sex can approach their differing fitness optima. In this inaugural study of sex bias in transcript structure across species, the presence of both male and female biased transcripts in the same gene provides evidence that alternative transcript structure, a mechanism of conflict resolution, may be widespread in Drosophila. We find that the sexually dimorphic transcripts we identify in this study tend to have distinct structural differences, such as alternative 5’ or 3’ exons, consistent with expectations of functional differentiation in the resolution of conflict.

Another prediction of the resolution of sexual conflict by sex differences in expression is that genes which show sex differences will also show reduced signatures of conflict, as has been observed in several studies^117–119^. While we do not have data specific to the focal species and strains used in our study, we can examine the correspondence of sex differences at the gene level to surveys of sexually antagonistic genes in *D. melanogaster*^118, 120^. Ruzicka et al. 2019^118^ found that sex-biased genes were depauperate among the set of genes identified as showing signatures of sexual antagonism – consistent with what we expect if the conflict is resolved by the evolution of sex bias. We do not see this trend in our data. Examining the correspondence of our gene level estimates of bias with genes identified as sexually antagonistic to a second study, Innocenti and Morrow 2010^120^, we find an increase in prevalence of sex bias over what is expected. This is the opposite of what we expect if the sex bias fully resolves sexual conflict. Additionally, genes identified as sexually antagonistic or under strong balancing selection in prior studies are among those showing sex differences in transcript usage contrary to this expectation (e.g. Fer2LCH, CG1637, Aldo; Figure 5B-C^118, 120, 121^). This indicates a more complex picture, with potential constraint on the full resolution of conflict^106^. Male and female bias in transcripts that are expressed from the same gene differ in transcript structure suggesting a response to sexual conflict, while the lack of negative correlation with evidence of sexual antagonism further suggests the conflict is not fully resolved.

The number of genes in a gene family and the number of alternative transcripts show negative correlation, as expected if they are alternative mechanisms of functional diversification^6, 107^. We find no association between sex bias and the relationship between these two mechanisms as might be expected if diversification along lines of sex drives the observed pattern with conflict resolved exclusively by one or the other mechanism. The focal tissues examined in this study generally show fewer sex differences in gene expression, for example relative to the gonad where duplication may be a common mechanism of generating sex-differential gene expression^122–124^. We do not see evidence for a role of duplication in the evolution of sex-biased or sex-specific gene expression in these tissues. Head tissues, including the brain, have high transcriptional diversity relative to other sets of tissues^125^. Multiple transcripts suggest greater possibility for evolutionary changes in response to context, due to relaxed constraint and enhanced capacitance for the evolution of functional differences and we find that the presence of multiple transcripts is associated with sex bias. This suggests that alternative splicing may be a key mechanism for the evolution of sex differences in transcript usage in these tissues, which could be explained by increased diversity, and variation, at the transcript level providing a substrate for selection.

### Sex bias in transcript usage evolves rapidly

Both sexual conflict and sexual selection can drive the rapid evolution of sex differences in gene expression^106, 126–129^. Consequently, sex differences are widespread, and the conservation/divergence of sex-biased expression has received considerable attention. However, whether this is due to gene level or transcript level regulation is unclear in the current literature (reviewed in^130, 131^). We see little evidence of uniform sex bias (all transcripts biased toward one sex). The lack of uniformity in sex bias among the observed transcripts of a gene suggests that the evolution of sex bias is not primarily due to differences in promoters that impact the overall rate of transcription of a gene, but rather due to mechanisms like alternative promoters and splicing that affect the production of specific transcripts in each sex. We also see little evidence for sex-limited transcripts suggesting that they are rare relative to quantitative differences between transcript structures present in both sexes. This implies that there is constraint on context specific transcription, either across sexes or across tissues and cell-types^132^. Examining sex differences in transcript orthologs, we find that between *D. melanogaster* and *D. simulans* there is greater conservation of female biased transcripts, relative to male biased transcripts, consistent with prior gene level studies and expectations based on patterns of sexual selection that result in more rapid evolution of male biased expression^133–135^. However, we do not observe this asymmetry for other species pairs and there are roughly similar male/female biased transcripts or more male-biased conserved transcripts (*D. simulans* and *D. serrata*).

### Sex bias in transcript usage can be conserved over long evolutionary distances

The set of highly conserved patterns of sex bias in transcript expression we identify among head expressed genes is intriguing. Over the greater distance, to *D. serrata*, shared patterns of sex bias may represent convergent evolution or turn-over, but some studies have demonstrated that multiple isoforms of a gene can be conserved over long periods of evolutionary time^13, 136, 137^. Among the most highly conserved genes and transcripts are those expressed from the Doublesex regulated yolk protein genes, and we identify an additional known Dsx target *Fmo2*^138^. We also find that genes encoding a set of interacting ribosomal proteins, and rna-binding protein Hoi-polloi (Hoip), are among the small set of genes showing conserved female-bias across *D. melanogaster*, *D. simulans* and *D. serrata*. Hoip is the *Drosophila* ortholog of the snRNP, small nuclear ribonucleoprotein 13 (hSNU13 ^139^). This protein has multiple regulatory functions, including in splicing and ribosomal small subunit biogenesis, and plays a role in neurodegeneration^139, 140^. One hypothesis that explains this pattern is that highly conserved sex differences in expression of these genes reflect regulatory interactions that underlie the ability to express high levels of proteins, like expression of yolk proteins in the fat body, in females. There are similar sets of interacting genes in males, related to glutamate and glutamine metabolism (*glutamate synthase*, *Glts*; *glutamine synthetase 2*, *Gs2*). While there are many possibilities, in this context this could reflect sex differences in behavior related to glutamatergic signaling^141, 142^. Interactions can constrain the resolution of sexual conflict or play a key role, and it is possible that once they have evolved sex differences among interacting gene products may similarly constrain the difference in expression and in gene function consistent with what we observe^105, 106^.

## Conclusions

The fiveSpecies annotation enables comparative evolutionary studies of alternative transcription in Drosophila. We gain the ability to compare expression of orthologous transcripts across species by amalgamating annotation information across species and integrating the inference of homologous relationships between transcripts into the annotation process - which our approach makes possible even in the face of variation in genome quality, apparent exon splits/fusions, and multiple alternative potential junctions. We present a suite of practical approaches to identify transcript orthologs and alternative transcript structure that can be replicated and applied to include other species of Drosophila and other genera. Expressed transcripts are both diverse and conserved, suggesting more biologically functional transcripts, for example in rare cell types, than current annotation suggests and highlighting a significant gap in our understanding of the evolution of gene function. Currently, functional annotation is at the gene level, perhaps due to prior limitations in parsing expression data into transcripts. The advent to long read technology is transformational-it enables us to empirically record transcript expression in a myriad of contexts. The next step is to associate transcript expression context, transcript structure and function as key components of annotation. Our adoption of a binary scoring of transcript structure relative to a gene model is completely generalizable and enables the computationally efficient comparison of transcript structure between contexts and could be used as a basis for association with functionally relevant sequence motifs. In order to do this systematically, the community needs to develop standards for recording context dependent expression of transcripts.

## Methods

### Liftover annotation

We performed a reciprocal liftover for each of the individual species’ annotations onto the genome co-ordinates of all species (Figure 1, Supplementary Figure 1). All mapping was conducted with minimap2 with parameters (minimap2 v2.24 parameters: -a, -x splice, --secondary=yes, -N 200, -p 0.9, -C 5) (Minimap2^143^). When the source transcript maps to an annotated transcript in the target species an edge is recorded. The relationships among individual species transcripts is tracked by considering each transcript a node and recording edges when (Figure 1A). We used a network graph to link homologous transcripts and consider any set of homologous transcripts with one transcript in each of the five species to be a potential orthologous set and describe the topology of the networks using standard topological definitions. Using the gene identifiers from the individual species we gathered all transcripts from the same gene within species and then used the associated species level gene identifiers at each node to identify genesets with linked components.

### Sample collection

*D. melanogaster* F1 progeny were collected from a cross between two laboratory strains (A7 T.7 (DSPR Founder Line – Tucson 14021-0231.7) females by B2 CA1 (DSPR Founder Line - BDSC 3846) males. *D. simulans*, F1 progeny were collected from a cross of MD106ts (DSSC 14021-0251.196) females by W501 males (DSSC 14021-0251.195). Flies were housed at Auburn in a 25 ⁰C incubator with a 12 hour light-dark cycle and reared on standardized medium (Bloomington Drosophila Stock Center cornmeal recipe). Virgin males and females were collected, aged in single sex vials, and flash frozen in liquid nitrogen under CO2 anesthesia within a 2-hour window. Each independent replicate consisted of a pool of 50 five-to-seven-day old animals. Vials were shipped to Florida on dry ice. *D. yakuba* and *D. santomea* were reared at a common temperature of 22.5-23.5C at UNC-Charlotte. *D. serrata* were collected at UC Davis. Adult males and females were collected within 5 days of eclosion. Flies were flash frozen in liquid nitrogen, and vortexed at high speed to remove heads. Samples were stored at −80 prior to extraction. Vials were shipped to Florida on dry ice. Fifteen heads were sampled from each replicate.

For each sample, mRNA was extracted and ONT barcoded libraries constructed according to the manufacturer’s protocol. *D. melanogaster* and *D. simulans* were pooled, *D. yakuba* and *D. santomea* were pooled and *D. serrata* samples were pooled with each pool sequenced on multiple PromethION flow cells (see Supplementary Methods). We generated updated Dorado basecalled reads from pod5 or fast5 files by converting them to pod5 formats (pod5 v 0.3.6) prior to basecalling by Dorado (v 0.5.2) (https://github.com/nanoporetech/dorado) using options --recursive --device “cuda:0,1” --kit-name SQK-PCB109 --trim none. Reads were demultiplexed using the demux mode of Dorado (v 0.5.2) with options --no-classify –emit-fastq. The resulting fastq files were processed using pychopper (v 2.7.1). We aligned the reads from each species to their own genomes using Minimap2 version 2.17 with the parameters “-12 -a -x splice –secondary=yes -N 200 -p 0.9 -C 5” (Minimap2^143^) and evaluated the quality of the experiment relative to the original annotation and the updated fivespecies annotation using SQANTI-reads^43^. The resulting read classifications relative to the fiveSpecies annotation were used to identify likely transcript fragments, and transcripts that match the annotation.

### Exon patterns

Gene models were estimated in TranD^23^ using the 1GTF gene mode and the current annotation from each species (Supp table: genome annotation). All transcripts were scored for their exon pattern relative to the gene model using the binary system described in TranD and the utility designed for this purpose. (Nanni 2023^23^ id_erp.py, Supplementary Methods). After removing likely transcript fragments identifies by SQANTI-reads^43^ transcripts are defined as having a pattern consistent with the annotation or novel relative to the annotation. We compare the structure of all transcript pairs using the binary indicators. Pairs with the same binary pattern were evaluated for alternative donors/acceptors and intron retention events. Pairs with different binary patterns were compared using set operations. An alternative start occurs when the minimum value in the two ordered sets differs and an alternative end when the maximum value differs. Exon skips occur when the difference between the two sets differ in least one member not on the boundary (pairwise_as_detection.py).

### Sex bias

Transcripts with enough coverage (more than 25 reads total and at least 5 reads in >50% of the samples for one sex) were evaluated for sex bias in *D. melanogaster*, *D. simulans* and *D. serrata* using a t-test. Transcripts with the same binary exon pattern, are also aggregated and tested for sex bias where possible (Supplementary Methods).

### Summarizing Data for Homologous Transcripts/Genesets

Each Unique Junction Chain (UJC) is linked to a unique component by the network graph. For each component in each species the sex bias results for the UJC(s) associated with that component were recorded. A single UJC in a component was considered simple and flagged. For components with multiple analyzed UJC if at least one was significant for sex bias the component was considered significant. Homologous transcripts were linked across species based on the component identifier. Each UJC is also linked to a gene and a geneset. Genesets may have more than one geneID in the case of overlapping genes (e.g. *Sxl*). Genesets are summarized from the associated UJC in the same way as components.

### Summarizing data for Genes

All the UJC associated with an individual GeneID are first summarized for the sex bias results based on whether the UJC was annotated, had an annotated exon pattern, was a likely fragment (ISM) or novel. Exon Region Pattern plus (ERPp) associated with a gene are summarized based on whether they were present in the annotation, likely representing a transcript fragment or novel. The number of UJC in the category, and the number statistically significant (for *D. melanogaster, D. simulans, D serrata*) is tallied. If at least one UJC or ERPp was significant for sex bias the gene was considered significant for sex bias.

### Comparing between species

The component ids and geneset ids are connected across species with the network graph. This allows one-to-one merges of results on these identifiers.

### Tests of Enrichment

We calculate the baseline frequency of sex bias for genes in *D. melanogaster* evaluated for sex bias using the categories, both, female, male, unbiased. For genes in a particular list, we compared the frequency of genes in the list in these four categories, to the baseline frequency in the data using a ChiSquare test with 3 degrees of freedom, and for the binary Biased/Unbiased with a ChiSquare test with 1 degree of freedom.

## Abbreviations

BP – Base Pair

ERP – Exon Region Pattern

ERPp – Exon Region Pattern plus

FSM – Full Splice Match

IRE – Iron Response Element

ISM – Incomplete Splice Match

UJC – Unique Junction Chain

## Acknowledgements

The Research Computing Team at the University of Florida was invaluable in their help, especially Olexander Moskalenko. The HiPerGator compute environment made all of this work possible. The University of Florida Genetics Institute and Department of Molecular Genetics and Microbiology and the Biology Department at Auburn, the Drosophila Genetics and Molecular Evolution conferences run by the Genetics Society of America and the Society of Molecular Biology and Evolution were all places where our colleagues listened to our ideas and offered invaluable advice and insights. We are grateful for the community of science.

## Data Availability

Fastq files are available at the SRA under 1) BioProject PRJNA1219500 for the *D. melanogaster* and *D. simulans* samples, 2) BioProject PRJNA1242287 for the *D. santomea* and *D. yakuba* samples and 3) BioProject PRJNA1220163 for the *D. serrata* samples.

All supplementary files are available at **Error! Hyperlink reference not valid.** Files include full annotation files (fiveSpecies_[species]_full_annotation_w_component.csv) and the associated fiveSpecies annotation files used to create the full annotation files (xscript_link files, GTF files, etc) and the datafiles (datafile_jxnHash_[species]_w_annoFlag_ERP_ESP_info_strCat_flagNovel_02amm.csv, datafile_erp_[species]_w_annoFlag_flagNovel_02amm.csv) along with relevant associated files.

Scripts and documentation are available on the GitHub page: https://github.com/McIntyre-Lab/papers/tree/master/bankole_sex_specific_splicing_2026

## Author’s Contributions

**AK, OB, RMG, RLR** collected all tissue. **AMM** extracted RNA, constructed libraries, and performed all bioinformatic analysis of the data from quality control to statistical testing. **KB** designed and executed the fiveSpecies annotation process and designed and executed assignment of exon patterns to reads. **AH** performed the duplicate vs alternative transcript analyses. **NK** evaluated the quality of the long read data and identified novel transcripts. **MG** designed and coded all transcript, gene and geneset summaries and made all the figures, developed and coded the algorithms for comparing transcript structure based on set theory designed and implemented the IGV viewer. **RLR** designed and interpreted the fiveSpecies annotation and analyzed results. **MK** contributed to Figures 1, 3 & 5. **LMM and RMG** designed the study, and analyzed the results. The manuscript was initially drafted by **LMM** and **RMG** with substantial input from **AK** and **RLR**. **AMM, KB** and **MG** provided documentation and scripts for the GitHub. The final manuscript was a joint effort of all authors.

## Ethics approval and consent to participate

Not applicable.

## Consent for publication

Not applicable.

## Supplementary Methods

### FiveSpecies annotation

Among the *Drosophila* with complete genomes, *D. melanogaster* is widely used in molecular and evolutionary genetics, with copius high throughput molecular datasets to facilitate annotation^1–4^. *D. melanogaster* is a classic model system with ∼35,000 transcript models, while *D. simulans D. yakuba*, *D. santomea* and *D. serrata*, have up to 40% fewer transcript models. However, it remains unclear whether even with this vast empirical data that non-model *Drosophila* might still hold clues to improve annotations due to their widespread use in studies of evolutionary genetics (Supplementary Table, transcript_model_counts.xlsx)^5, 6^. We postulated that transcript models would be conserved across species, and that existing annotation in the individual species could be leveraged to improve the annotation across all five species. We developed a ‘liftover’ strategy^7–9^ based on the success of our recent study in the closely related species *D. melanogaster* and *D. simulans*^10^. We lifted annotations between species by mapping FASTA files from each of the five transcriptome sources (Supplementary Table, genome_sourcefiles.xlsx) onto the other four target genomes.

To minimize discrepancies between annotated positions on the individual species genome co-ordinates and positions derived from mapping reads from the data, we also mapped the FASTA files from the source to their own reference genomes. All mapping was conducted with minimap2 with parameters (minimap2 v2.24 parameters: -a, -x splice, --secondary=yes, -N 200, -p 0.9, -C 5) (Minimap2 Li 2018). Annotated transcripts that are unmapped, supplementary, or have a map quality equal to 0 were tracked. To mirror the data handling for the experiment, gene identifiers were assigned to transcripts via GFFcompare^11^, and any discrepancies with annotated gene identifiers noted (cross_species_link_files/annotation_2_genome_compare_transcript_gene_assignment.csv, annotation_2_genome_flag_ambig_gene.csv). In *D. melanogaster* and *D. simulans* some genes had multiple identifiers, we selected the first alphabetical identifier and tracked the synonyms (fiveSpecies_annotations/fiveSpecies_2_genome_anno_files). We summarized the structure of the individual species mapped transcript models using the 5’ to 3’ list of genomic coordinates that indicate the exon/intron boundaries, which we refer to as the junction chain^10^. Junction chains vary in length based on the number of exons, for computational efficiency and data management consistency, we use a 64-character hexadecimal hash, the junction hash (jxnHash), based on SHA-256^12^. This encoding does not alter the underlying genetic structures of isoforms but rather offers a uniform data structure to encode this genetic information in high throughput analysis. Transcript models with identical mapped junctions (*i.e.* identical splicing breakpoints) will have the same unique junction chain (UJC) and the same junction hash. A GTF file with the representative transcript model for each UJC consisting of the 5’ and 3’ most ends is created with the naming convention annotation_2_genome_ujc.gtf. The relationships between the 5’ and 3’ ends from the annotated transcript models, and the UJC are tracked (Figure 1; annotation_2_genome_ujc_xscript_link.csv). The representative UJC are used to extract a transcript FastA file from each of the five source genomes. Each source FASTA is mapped onto the remaining four target genomes with minimap2 (minimap2 v2.24 parameters: -a, -x splice, --secondary=yes, -N 200, -p 0.9, -C 5) (/submission/supplementary/fiveSpecies_annotations/fiveSpecies_2_genome_anno_files annotation_2_native genome_ujc_2_target genome_noGeneID_ujc_xscript_link.csv) and assigned a junction hash based on the target species co-ordinates representing the orthologous match for a given transcript in the cross-species mapping. The resulting set of UJC for each of the five target species are summarized using the junction string/junction hash strategy described above (annotation_2_genome_ujc_xscript_link.csv) in order to simplify the computational effort in tracking unique identifiers on each set of genome co-ordinates.

Orthologous isoforms from different source species may map to a common position in the target genome, or, given the sequence divergence, may map to slightly different positions. If the mapping is not-identical these isoforms will not share UJC, resulting in complex relationships across isoforms from the different source genomes. The relationships among isoforms can be tracked by considering each UJC as a node and mapping all of the fiveSpecies source UJC to the other four target genome co-ordinates, recording an edge if the mapped UJC matches a node in the target genome (Figure 1A). The network is a set of components linking the species nodes via edges (edges.csv, component_map.csv). Mapping across species with this level of sequence divergence can lead to complex topological relationships. We record the number of species linked by each component. If the nodes in the source and target genomes are connected by reciprocal edges in all five species this topology is referred to as a k5. If nodes are connected by reciprocal edges in 4 of the 5 species, and the fifth species is connected to a single node this is referred to as a pendant. We tracked these simple topological structures.

For all of the components we classify the origin of the transcript model and the number of species with an annotated transcript in the component using the phylogeny of the species (Supplementary Figure 1) where we label the following relationships, all five species (‘all five’), all but *D. serrata* (‘the four’), and the branches (‘melsim’, ‘yaksan’), each species ‘only’.

### Comparing the Lift Over to Orthologs

We generated transcript annotations across the entire genome generating a complete map of isoforms across the five *Drosophila* species. These transcripts were identified in the absence of gene location data so that each transcript is placed without bias from prior gene annotations that might exclude portions of transcript models. Transcript models and their related UJC identifiers are then assigned to genes using GFF compare on each species co-ordinates and genes are linked across species based on the linking of the transcript models. If as a function of liftover a transcripts is located in a region not previously annotated as a gene, GFFcompare assigns an arbitrary identifier, we updated this identifier in the annotation files with a “GFF5_” prefix in order to make clear that the gene was novel to the target species. The resulting set of UJC and their corresponding gene identifiers on each species co-ordinates is the fiveSpecies annotation for each set of genome co-ordinates. Using the GFF compare assigned geneIDs we linked the components across species into genesets. We then compared genes in the genesets to known lists of orthologs. For *D. melanogaster* and *D. simulans* we used the Flybase precomputed file (FB2022_01 from ORthoDB) to identify orthologs and for *D. yakuba, D. santomea* and *D. serrata* we downloaded the orthologs from NCBI ortholog table in March 2025 (sex_specific_splicing/adp_ortholog_lists).

### Exon Region Patterns

We summarize the complete exon content for each gene across all potential isoforms identified in the Five Species Annotations. This list of Exon Regions is typically greater than the exon content of any single isoform as it represents the combined set of exons across every alternatively spliced mRNA. Co-ordinates for exon regions were defined as the 5’ most acceptor and 3’ donor. We track donor/acceptor and 5’ / 3’ variation by tracking exon segments^10, 13^. We note that the donor/acceptors here are defined by mapping to facilitate the comparisons to the data and across species and they may differ from the annotated splice junctions. We introduce a new notation for individual transcript models based on their shared substrate of potential exon regions across the gene to facilitate comparisons (Figure 3). For each transcript model a value of 1 is assigned when the transcript model overlaps with an annotated exon region, and a 0 otherwise. For example, +_1-1-1-1 indicates a transcript from a gene annotated on the positive strand with 4 exon regions that contains all of the annotated regions, while +_1-0-1-1 indicates a transcript model that does not contain an exon in the second region. This binary summary of the exon regions included in the transcript model is called an exon region pattern (ERP) (fiveSpecies_annotations/fiveSpecies_2_genome_anno_files fiveSpecies_2_genome_ujc_er_vs_fiveSpecies_2_genome_ujc_infoERP.csv). Each network component with a single annotated exon pattern per species represents a single orthologous transcript and was considered a ‘simple’ component. Genes with a single simple component were classified as single transcript in the annotation. Genes with a single exon were categorized as mono-exon. All other genes were designated at multi-transcript in the annotation.

### Five Species Annotations

Representing these comprehensive data and the full diversity of alternatively spliced RNA sequences is challenging in a single file format. To fully capture the representative information from the transcriptome, we offer five files for each species that identify transcript models for the entire transcriptome. The resulting annotation files for each species contain, a GTF file for representative UJC; A GTF and a csv file for the exon regions for each gene; A GTF and a csv file for the exon segments for each gene; a csv file unique on the junction hash with a set of binary indicators linking the UJC to the source annotations from all five species, and a component map that links the species. The information for each file format is available in the supplemental files. Users wishing to generate the complete CDS sequences for the species should use the GTF file for the UJCs and a script (make_refernce_ujc.py will create a fasta file of the representative UJCs). When the number of exon regions is the same between species, we are also able to computationally compare ERP and determine if we have evidence for conservation of particular splicing patterns in a straightforward way. There are a number of reasons why the number of exon regions may differ between species pairs including potential exon splits/fusions and these can be identified. There may be real evolutionary differences that drive intron formation. There may also be artifacts due to mapping nucleotide sequences across the deep evolutionary timescale, and there may be errors and inconsistencies in the genome assemblies. The network of components combined with the ERP identify potential splits/fusions for all of the linked transcript models at scale.

### Drosophila samples, library construction and sequencing

We generated the Five Species Annotations to facilitate transcriptome analysis in biological data. To identify sex specific expression of alternatively spliced transcripts, we collected long read sequences from each of the five species. A total of 28 samples were sequenced using Oxford Nanopore Technology (ONT). For *D. melanogaster*, we sequenced 12 samples (2 sexes * 6 replicates) each consisting of 15 F1 heads from a cross between DSPR founder lines A7 T.7 ‘TAIW’ (Tucson 14021-0231.7) and B2 CA1 (BDSC 3846). For *D. simulans*, we sequenced a total of 12 samples (2 sexes * 6 replicates) consisting of 15 F1 heads from a cross between MD106ts (DSSC 14021-0251.196) and W501 (DSSC 14021-0251.195). For *D. serrata*, we sequenced a total of 12 samples (2 sexes * 6 replicates) consisting of 15 F1 heads from a cross between dark 681.5 and light 104. For *D. santomea*, we sequenced 2 samples (2 sexes) consisting of ∼45 heads from STO-CAGO 1402-3, Drosophila Species Stock Center 14021-0271.01. For *D. yakuba* we sequenced 2 samples (2 sexes) consisting of ∼45 heads from Tai18E2, Drosophila Species Stock Center 14021-0261.01. All samples were flash frozen, freeze-dried prior to isolating head tissue.

ONT libraries were constructed using the ONT PCR-cDNA Barcoding Kit (SQK-PCB109) starting with polyA mRNA according to the manufacturer’s protocol. The *D. melanogaster* and *D. simulan*s sample replicates 1-3 were processed and libraries constructed as a set of samples for pooling and sequencing. *D. melanogaster* and *D. simulans* sample replicates 4-6 were processed and libraries constructed as a separate set of samples for pooling and sequencing. The *D. santomea* and *D. yakuba* samples were processed and libraries constructed together as a set of samples for pooling and sequencing. The *D. serrata* samples were processed and libraries constructed as a final set of samples for pooling and sequencing. Within each set of samples, libraries were pooled to a total of 100 fmol and run on a MinION Mk1c with real-time basecalling and demultiplexing (Guppy v6.1.5, MinKNOW v22.05.8). For *D. melanogaster*, *D. simulans* and *D. serrata* samples, we used the MinION read counts to repool the libraries prior to obtaining additional sequencing data on the ONT PromethION (Guppy v5.1.13, MinKNOW v23.04.5) at the University of Florida Interdisciplinary Center for Biotechnology Research (ICBR, RRID:SCR_019152). Technical replicates (TRs) are defined as the same library run on different ONT flow cells (MinION or PromethION). TRs 1-3 were run on the MinION and TRs 4-6 were run on the PromethION.

The *D. melanogaster*, *D. simulans* and *D. serrata* data samples were stored in fast5 and pod5 format and as .fastq files. We generated updated Dorado basecalled reads directly from pod5 files or from pod5 files converted from pod5 formats (pod5 v 0.3.6) prior to basecalling by Dorado (v 0.5.2) (https://github.com/nanoporetech/dorado) using options --recursive --device “cuda:0,1” --kit-name SQK-PCB109 --trim none. Reads were demultiplexed using the demux mode of Dorado (v 0.5.2) with options --no-classify –emit-fastq. The resulting fastq files were processed using pychopper (v 2.7.1). *D. santomea* and *D. yakub*a data samples were stored as .fastq files only. The fastq files for the *D. santomea* and *D. yakuba* files were processed using pychopper (v 2.7.1).

### Read Mapping

We aligned reads from each species to their own genomes using Minimap2 version 2.17 with the parameters “-12 -a -x splice –secondary=yes -N 200 -p 0.9 -C 5” (Minimap2^14^. We converted sam files to gtf and assigned genes from the Five-Species annotation for the genome to each of the files of aligned reads using gffcompare^11^. Reads assigned to super-loci labeled as “XLOC” genes were tracked. For each gene, technical replicates were combined to identify UJC. If a UJC was assigned to different genes in different replicates, these are identified and separated from UJCs unambiguously assigned to specific genes.

The number of reads per UJC per sample was tabulated directly from the alignment files and the number of reads per gene in each sample was tabulated by summing across all un-ambiguously assigned UJC for that particular gene. All UJC that mapped to an annotated gene region were assigned an ERP and an ESP using the gene model summary from the Five-Species annotation for that species and UJC sharing an ERP were summed to count the number of reads per ERP per sample.

### Quality control

The proportion of the total number of reads that mapped to the genome, the length of each read and evidence for artifacts due to library preparation and sequencing were determined for each sample. Mapped reads were evaluated relative to the native species annotation and the Five-Species annotation. We tabulated: the proportion of mapped reads in genes; the proportion of reads where all of their mapped junctions exactly match an annotated transcript, or a full-splice match (FSM); the proportion of reads with a contiguous subset of mapped junctions or incomplete splice match (ISM); the proportion of mapped reads whose donor/acceptor sites are all in the annotation but not present in the observed combination, novel-in-catalog (NIC); and the proportion of mapped reads with at least one donor/acceptor that is novel, novel-not-in-catalog (NNC), using the tool SQANTI-reads^15, 16^. Additional classifications refer to reads not mapped in annotated genes. The difference between the native species annotation and the Five-Species annotation in terms of the proportion of reads classified in annotated genes and matching transcript annotation compared to the proportion without transcript annotations are used to evaluate the liftover strategy.

### Quantification of expression

An annotated transcript model (UJC) had empirical support when at least one read supported (FSM) that model. Note that reads from the data may have introns (noted with a binary indicator variable flagIR) and/or novel exons not present in in the Five-Species annotation (noted with a binary indicator variable flagDataOnlyExon). For each species, the distribution of reads per gene for each sample was used to calculate a normalization factor to account for the differences in the number of mapped reads in genes using the sample third quartile^17, 18^.

### Identification of novel transcripts

We used the ISM (incomplete splice match) label from SQANTI-reads to identify transcript fragments, and FSM to identify annotated transcripts in the data. If UJC had an ERP consistent with the annotation this was also flagged and the remaining UJC were considered to represent novel transcripts.

### Sex limited transcripts and Sex bias

We classified UJCs as sex limited if there were no reads in one sex for any replicate and in the other sex there were at least 50 reads. For *D. melanogaster, D. simulans* and *D. serrata*, a UJC was considered statistically analyzable for differences in abundance if there were at least 25 reads for the gene and at least 5 reads in 50% of the replicates for that UJC from either sex. Reps 4, 5 and 6 were used for the statistical analysis of differential expression. Short read RNA-seq studies have reported differences in variance between conditions, we tested for heteroscedasticity between males and females^19^ and for differences between the means allowing for heteroscedasticity^20, 21^ and we also calculated a test-statistic which assumed pooled variance^22^. We did not see evidence for widescale heteroscedasticity and used the test with the pooled variance. We used a nominal threshold of p = 0.05 to minimize type II errors in the comparisons across species. For *D. yakuba* and *D. santomea* we considered UJC potentially sex-biased if they were in a gene that was expressed and had 10 times or more reads in one sex. We compared results between *D. melanogaster* and each of the other four species when there was a linked UJC from the Five-Species annotation. If UJC shared the same ERP, we also tested the ERP for sex bias. A gene was considered to have evidence for sex bias if there was at least one test with statistical evidence for sex bias.

## Supplementary Figures and Legends

**Supplementary Figure 1.**
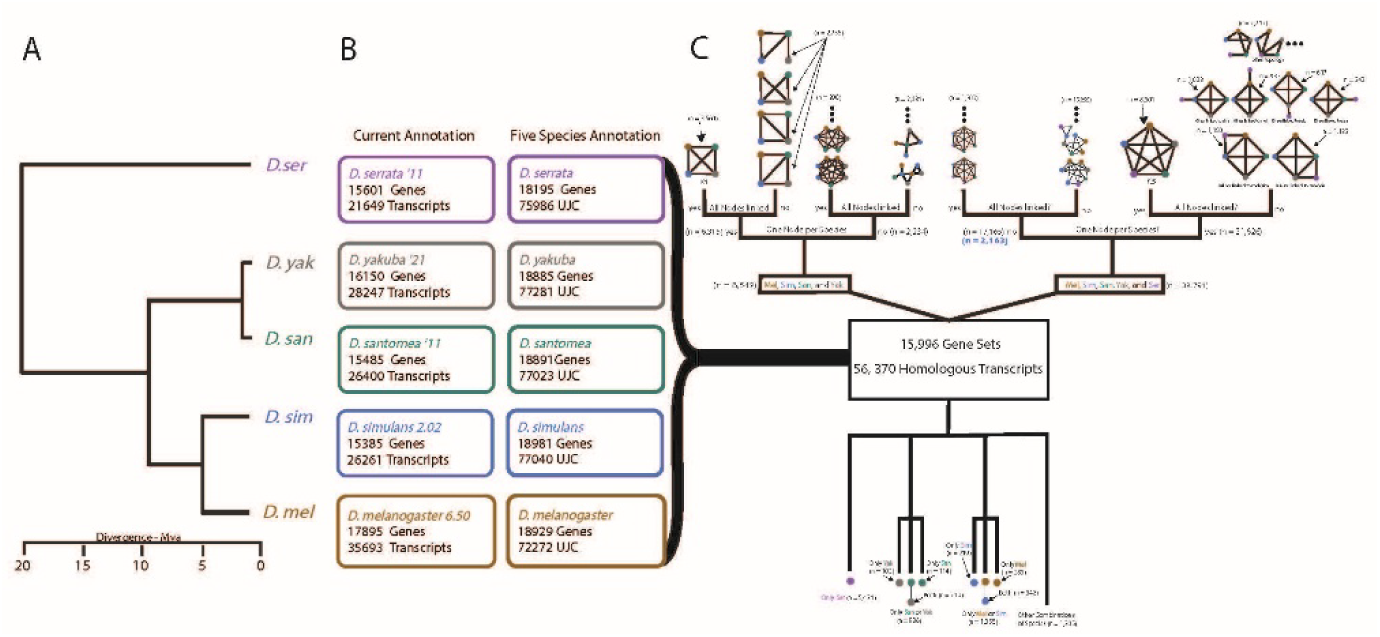
The fiveSpecies Annotation. Panel A) The phylogeny of *D. melanogaster* (*D. mel)*, *D. simulans* (*D. sim*), D. yakuba (*D. yak*), *D. santomea* (*D. san*) and *D. serrata* (*D. ser*). Panel B) The UJC counts for the mapped current annotation for each of these species (Supplementary Table: Genome annotation) and for in the fiveSpecies annotation after reciprocal liftover. The annotation files are in the supplementary data files fiveSpecies_<SPECIES>_full_annotation.csv. There are a total of 379,602 nodes in the network (75,986+77,281+77,023+77,040+72,272). Panel C) Topological description of the components from the network graph of edges between these nodes. There are 1,419,706 edges. The top right describes the topologies for components that contain nodes from all five species (38,791). 56% (21,626) contain a single node per species and are considered orthologs. Multiple nodes are likely a result of variation from mapping across species. If the data support a single node per species in these components we also consider them orthologs (2,163) for a total of 23,789 orhtologs. The second largest set of sub-graphs is for components with nodes in *D. melanogaster* (*D. mel)*, *D. simulans* (*D. sim*), D. yakuba (*D. yak*), *D. santomea* (*D. san*). Together comprising 84% of all of the annotation. There are some nodes that are unlinked to other species, these are relatively few with the exception of the outgroup D. serrata. There are some lineage specific homologous transcripts and some homologies between species that do not neatly match the phylogenetic expectations for the lineages studied here. In each of the individual fiveSpecies annotation files there is a unique component identifier, and the component_map_summary.csv table is a list of all the individual species nodes linked to the component identifiers. Some of the network structures we observe are common topological structures and these are recorded in the component_summary.csv.

**Supplementary Figure 2.**
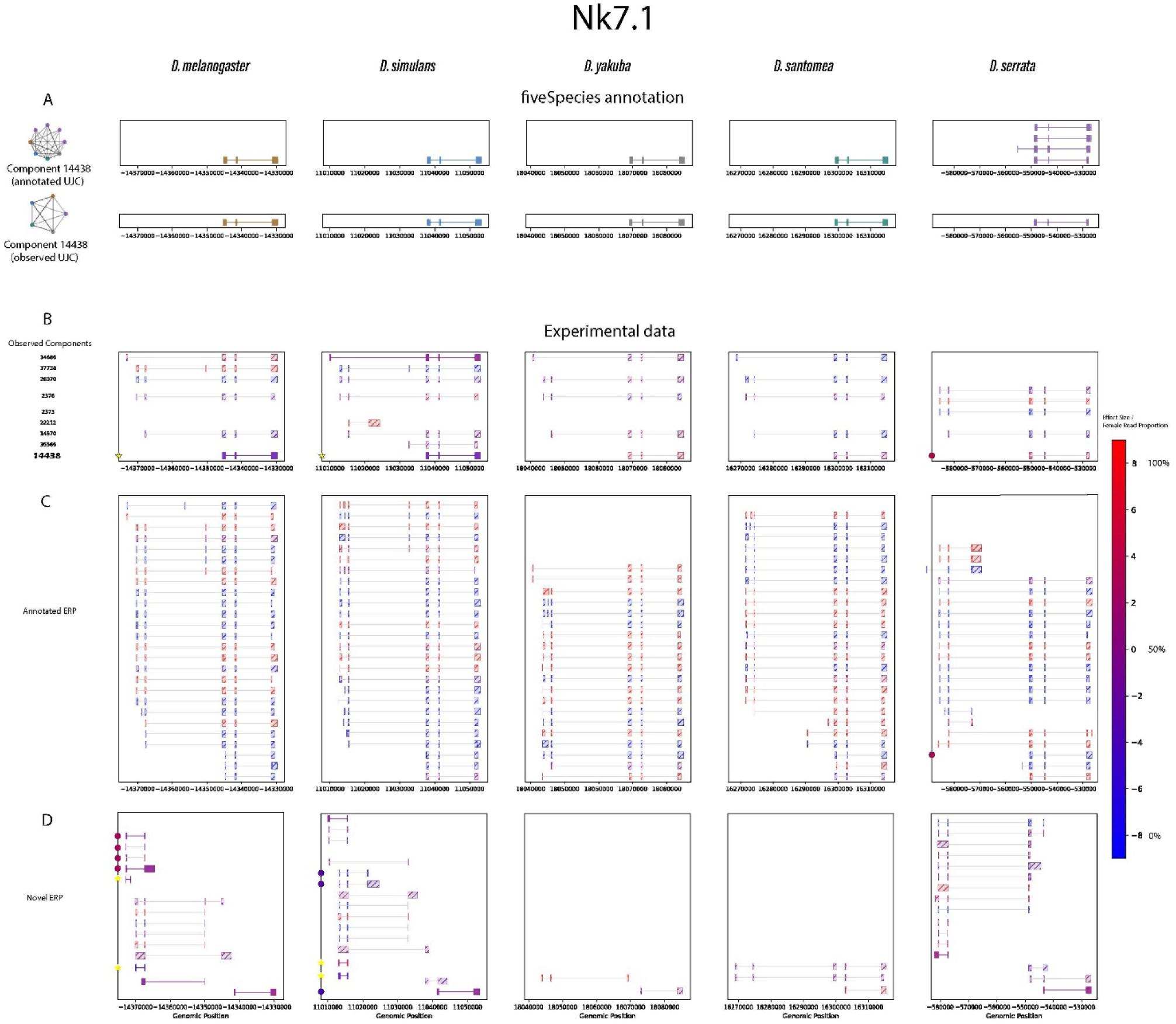
*Nk7.1*. Panel A) On the top row are annotated transcripts in component 14438. There are 4 annotated transcripts in *D. serrata*, and one in each of the other four species. On the top left is the network graph showing the edges between the annotated transcripts. The second row shows the observed transcripts for this component, and the network graph with only the observed nodes depicted. The four annotated transcripts in *D. serrata* differ only slightly and the long read data from *D. serrata* identifies one of the four as expressed. Panel B) Transcripts in the data. The top panel are annotated transcripts. The left most box indicates the corresponding componentID and contains all annotated transcripts with at least 1 read, the dark shading indicates there is sufficient data to analyze the transcript statistically for sex bias. The shading is on a heat map gradient by the effect size (right bar) with red indicating female bias and blue indicating male bias. Statistical significance (p<0.05) for the transcript is indicated by a yellow star. The filled circle indicates the set of transcripts that share the exon pattern are significant for sex-bias with the shading reflecting the effect size. Transcript 14438 is an alternate 5’ start to the compared to the other transcripts and was lifted onto D. melanogaster. The transcript ortholog is sex biased toward males in *D. melanogaster* and *D. simulans* but transcripts with the orthologous exon pattern are biased towards females in *D. serrata*. Panel C) Novel transcripts with annotated ERP. Transcripts with at least 10 reads are depicted. The filled circle indicates the set of transcripts that share the exon pattern are significant for sex-bias with the shading reflecting the effect size. Panel D) Novel transcripts with novel ERP. Transcripts with at least 50 reads are depicted. A potentially novel conserved transcript observed in *D. melanogaster, D. simulans* has significant male bias.

**Supplementary figure 3.**
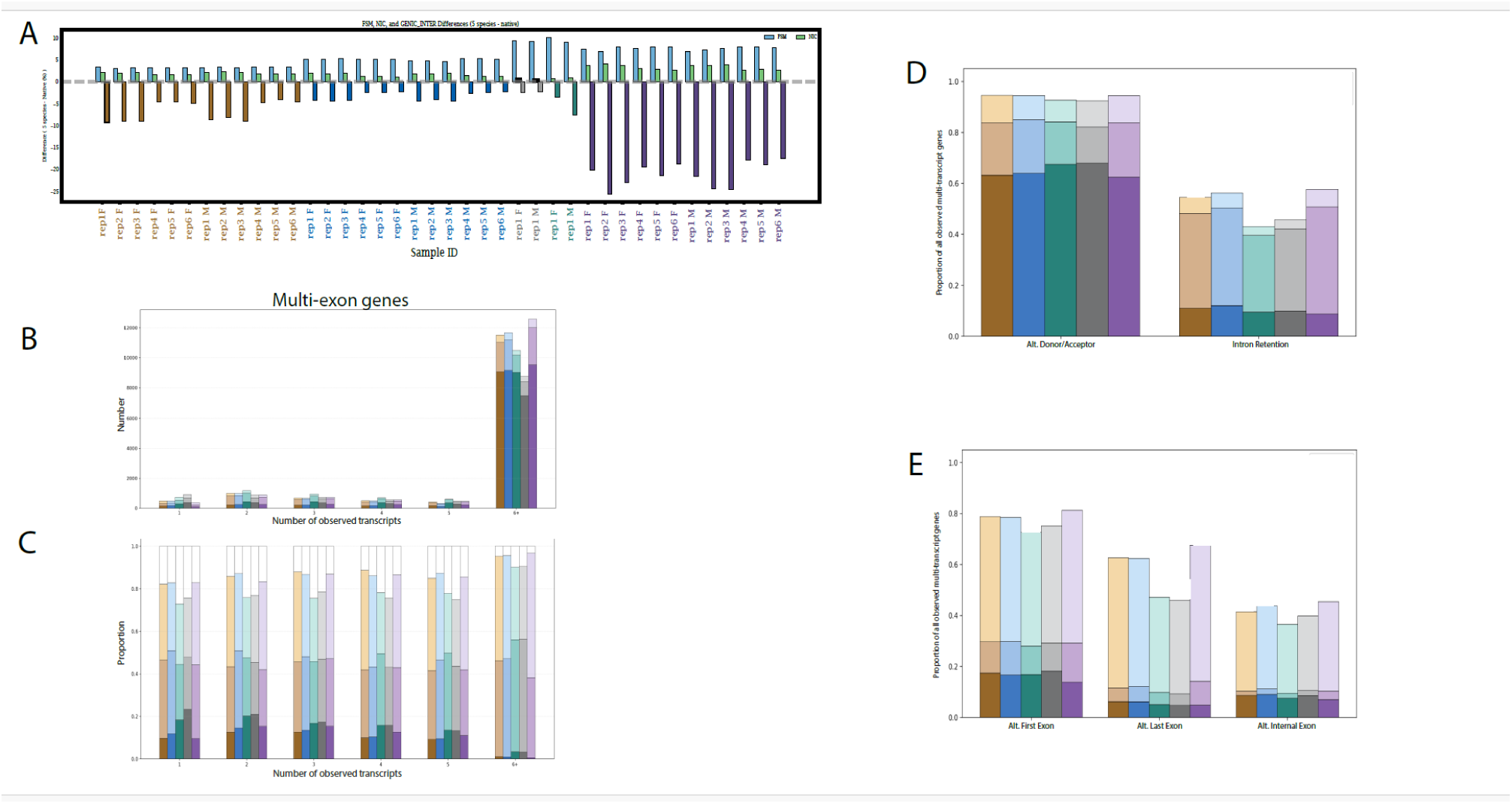
The fiveSpecies annotation. Panel A) The five species annotation compared to the current annotation. A bar plot for each of the biological samples, The Y axis is the difference in the proportion of reads assigned to each structural category between the fiveSpecies annotation and the individual species annotation. A positive difference reflects an increase in the proportion in the fiveSpecies annotation and a negative difference reflects a decrease in the proportion in the fiveSpecies. Blue bars represent reads whose junctions exactly match the annotation (full-splice-match FSM), green bars are reads consistent with annotated exon boundaries (novel-in-catalog NIC), reads that are genic-intergenic (negative) are colored by the species (*D. melanogaster*-gold, *D. simulans*-blue, *D. yakuba* gray*, D. santomea* green and *D. serrata* purple) Panel B) The Y axis is the number of observed multi-exon genes in *D. melanogaster* (gold, n=14,626), *D. simulans* (blue, n=14,380), *D. yakuba* (gray, n=12,338), *D. santomea* (green, n=14,740) and *D. serrata* (purple, n=15,648). The X axis is the number of observed transcripts. The transcripts that are annotated have dark shading, have an exon pattern consistent with the annotation are medium shading, or novel light shading. All UJC with at least 1 read that are not likely transcript fragments (ISM) are included. Panel C) The X axis is the number of observed transcripts. The Y axis is the proportion of transcripts that are annotated have dark shading, have an exon pattern consistent with the annotation are medium shading, or novel light shading. UJC with at least 1 read that are not likely transcript fragments (ISM) are included. Panel D) The denominator is the number of multi-exon genes. All pairs of transcripts that share the same exon pattern are compared. The bar indicates the proportion of genes with at least one pair with an alternative donor/acceptor and/or intron retention. The categories are not mutually exclusive. Dark shading indicates at least one transcript is annotated, medium shading indicates at least one transcript has an exon pattern consistent with the annotation and light shading indicates all transcripts that share the exon region are novel. UJC with at least 1 read that are not likely transcript fragments (ISM) are included. Panel E) The denominator is the number of multi-exon genes. All pairs of transcripts that differ in their exon pattern are compared. The bar indicates the proportion of genes with at least one pair with an alternative 5’ exon, alternative 3’ exon and/or an exon skip. The categories are not mutually exclusive. Dark shading indicates at least one transcript is annotated, medium shading indicates at least one transcript has an exon pattern consistent with the annotation and light shading indicates all transcripts that share the exon region are novel. All UJC with at least 1 read that are not likely transcript fragments (ISM) are included.

**Supplementary Figure 4.**
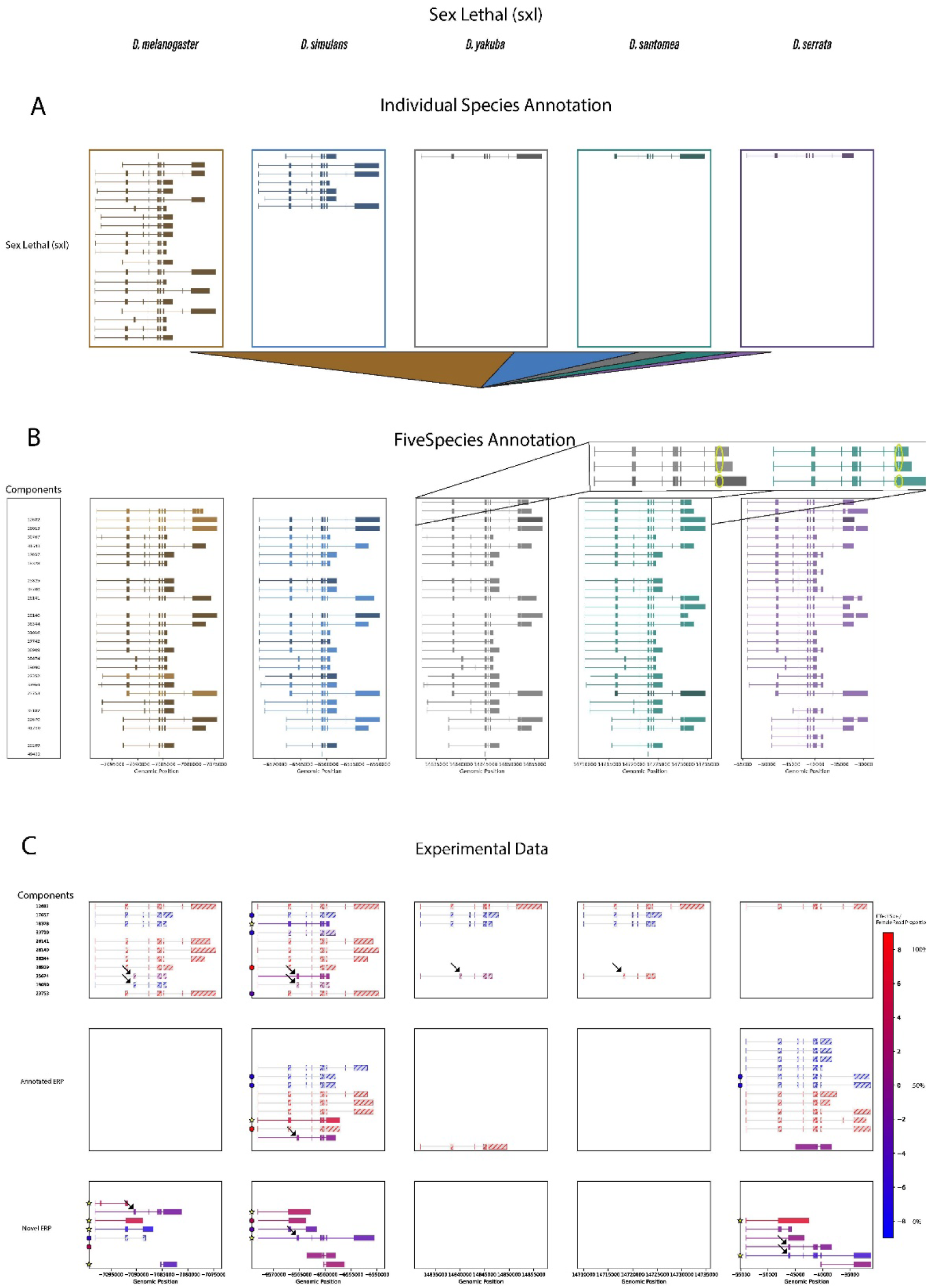
*Sxl*. Panel A) Individual species annotation for the *Sxl* gene. *D melanogaster* (n=20, gold), *D. simulans* (n=7, blue), *D. yakuba* (n=1 gray), *D. santomea* (n=1 green) and *D. serrata* (n=1 purple). Panel B) The fiveSpecies annotation after liftover and identifying homologous transcripts (components) using network graphs. The Component identifiers are listed in the left box, and transcripts with support from the empirical data are shaded. The zoom out shows component 12682 with a small intron present in the top 2 transcripts but absent in bottom transcript. The bottom transcript is supported by the data. Panel C) The experimental data for genesetid 1446 (*Sxl* and the overlapping lncRNA *D. mel* FBgn 0263563) for each of the five species. The top row shows annotated transcripts with at least 1 read, the second row shows transcripts with annotated ERP and at least 10 reads, the bottom row shows novel transcripts with at least 50 reads. Black arrows label exon Z. The shading is on a heat map gradient by the effect size (right bar) with red indicating female bias and blue indicating male bias. Statistical significance (p<0.05) for the transcript is indicated by a yellow star. The filled circle indicates the set of transcripts that share the exon pattern are significant for sex-bias with the shading reflecting the effect size.

**Supplementary Figure 5.**
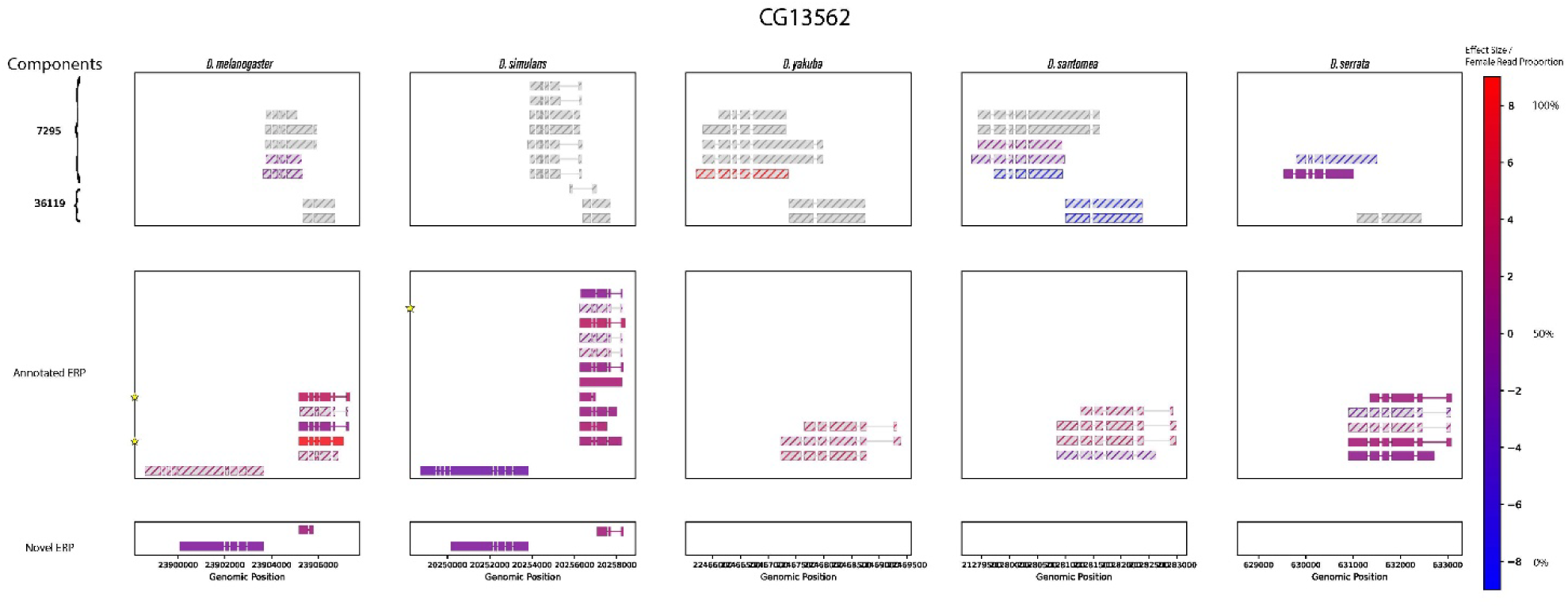
*CG13562*. The under annotated gene with a potentially novel conserved transcript. The fiveSpecies annotation *D melanogaster* (gold), *D. simulans* (blue), *D. yakuba* (gray), *D. santomea* (green) and *D. serrata* (purple). The top row shows annotated transcripts with at least 1 read, the second row shows transcripts with annotated ERP and at least 10 reads, the bottom row shows novel transcripts with at least 50 reads. Transcripts with some data are shaded. Transcripts analyzable for sex bias are shaded based on the heat map gradient by the effect size (right bar) with red indicating female bias and blue indicating male bias. Statistical significance (p < 0.05) for the transcript is indicated by a yellow star. The filled circle indicates the set of transcripts that share the exon pattern are significant for sex-bias with the shading reflecting the effect size.

**Supplementary Figure 6.**
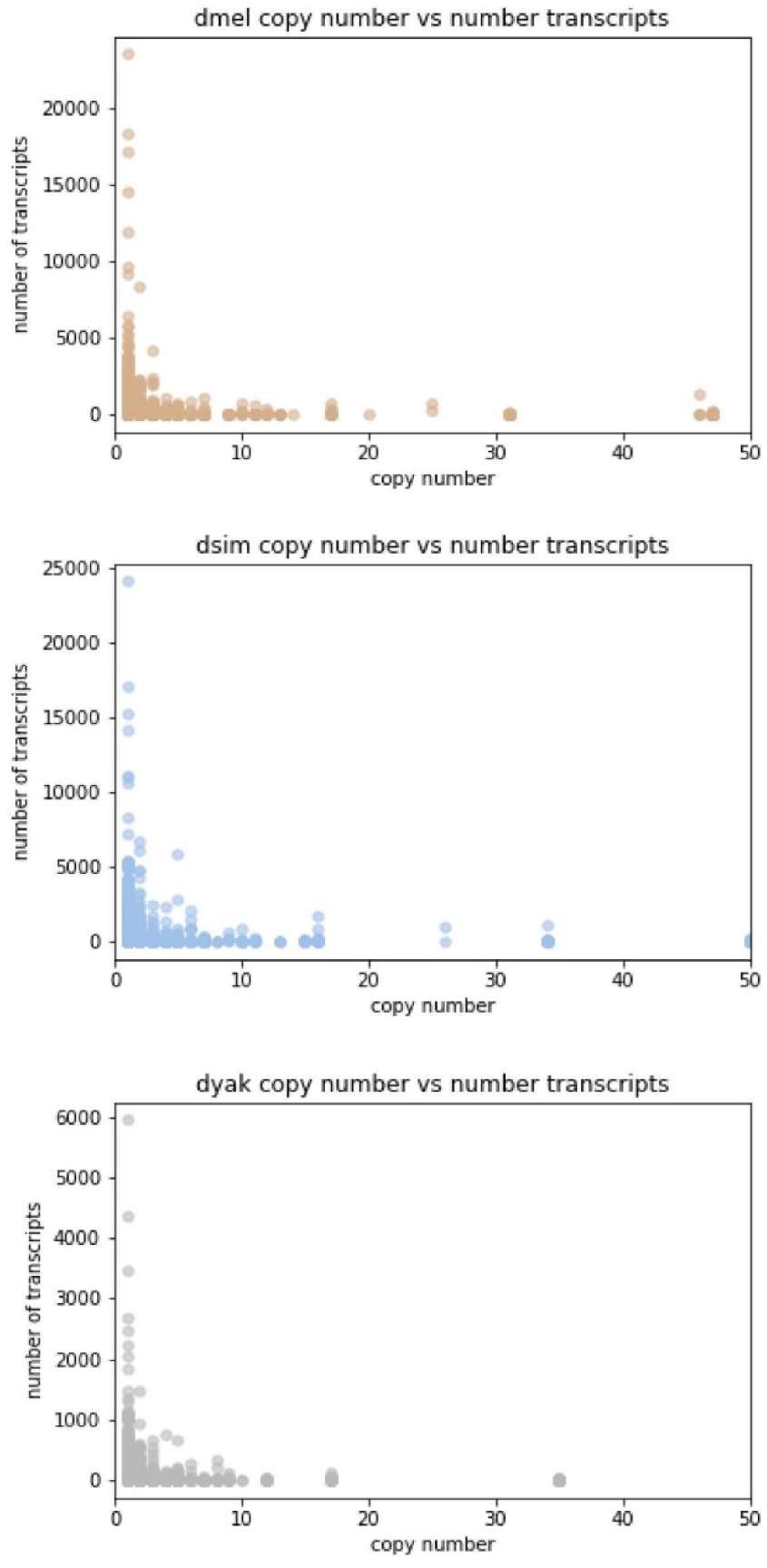
Alternative splicing and gene duplication. The Y axes are the number of transcripts with distinct exon patterns (not counting likely transcript fragments) and the X axes are the copy number. Each circle is a gene. top) *D. melanogaster*, middle) *D. simulans,* and bottom) *D. yakuba.* The X-axis is truncated there is a gene cluster at ∼200 copy’s in each of the three species.

**Supplementary Figure 7.**
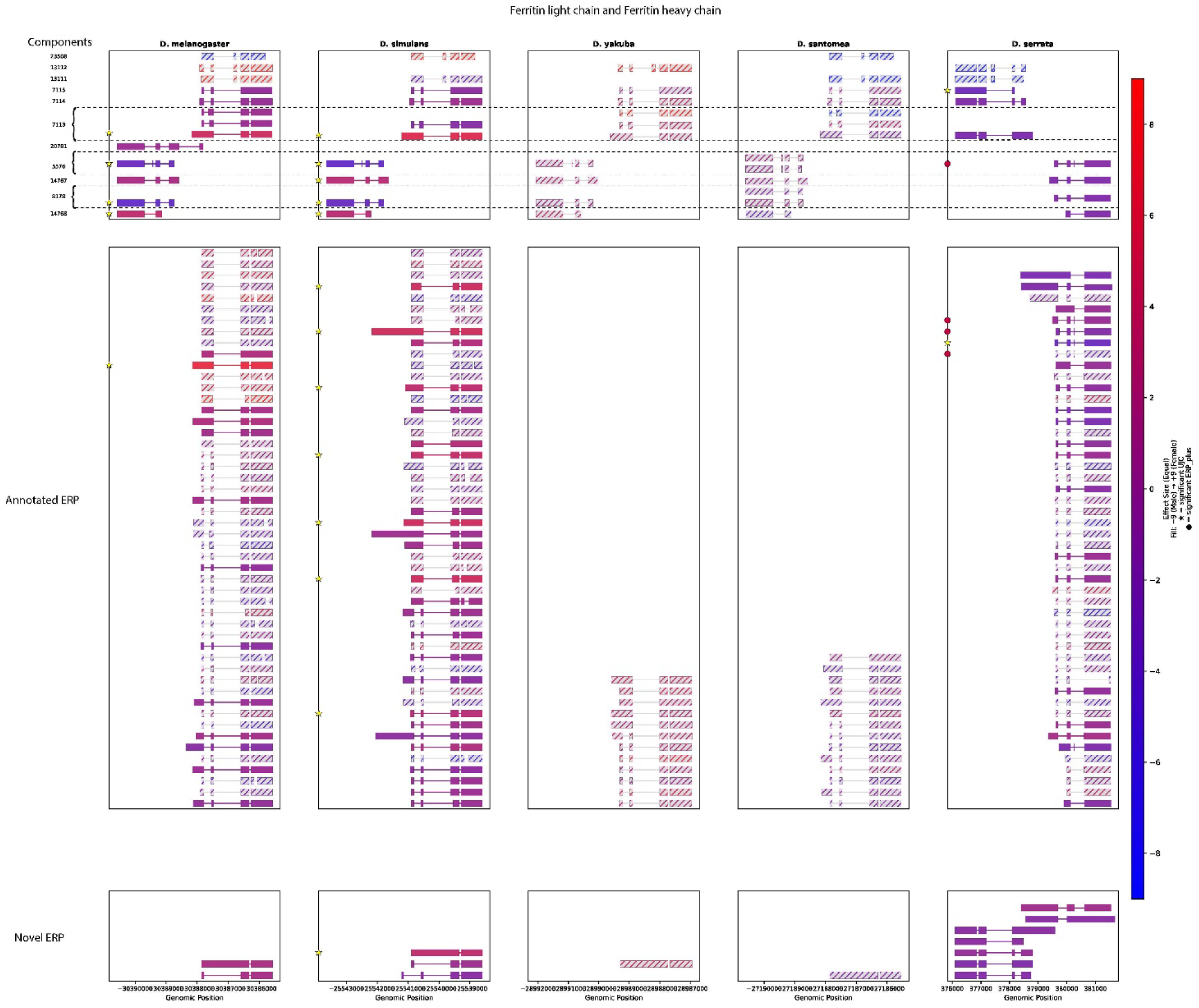
The *Fer1HCH* locus. The experimental data for each of the five species. The top row shows annotated transcripts with at least 1 read, the second row shows transcripts with annotated ERP and at least 10 reads, the bottom row shows novel transcripts with at least 50 reads. The shading is on a heat map gradient by the effect size (right bar) with red indicating female bias and blue indicating male bias. Statistical significance (p<0.05) for the transcript is indicated by a yellow star. The filled circle indicates the set of transcripts that share the exon pattern are significant for sex-bias with the shading reflecting the effect size.

**Supplementary Figure 8.**
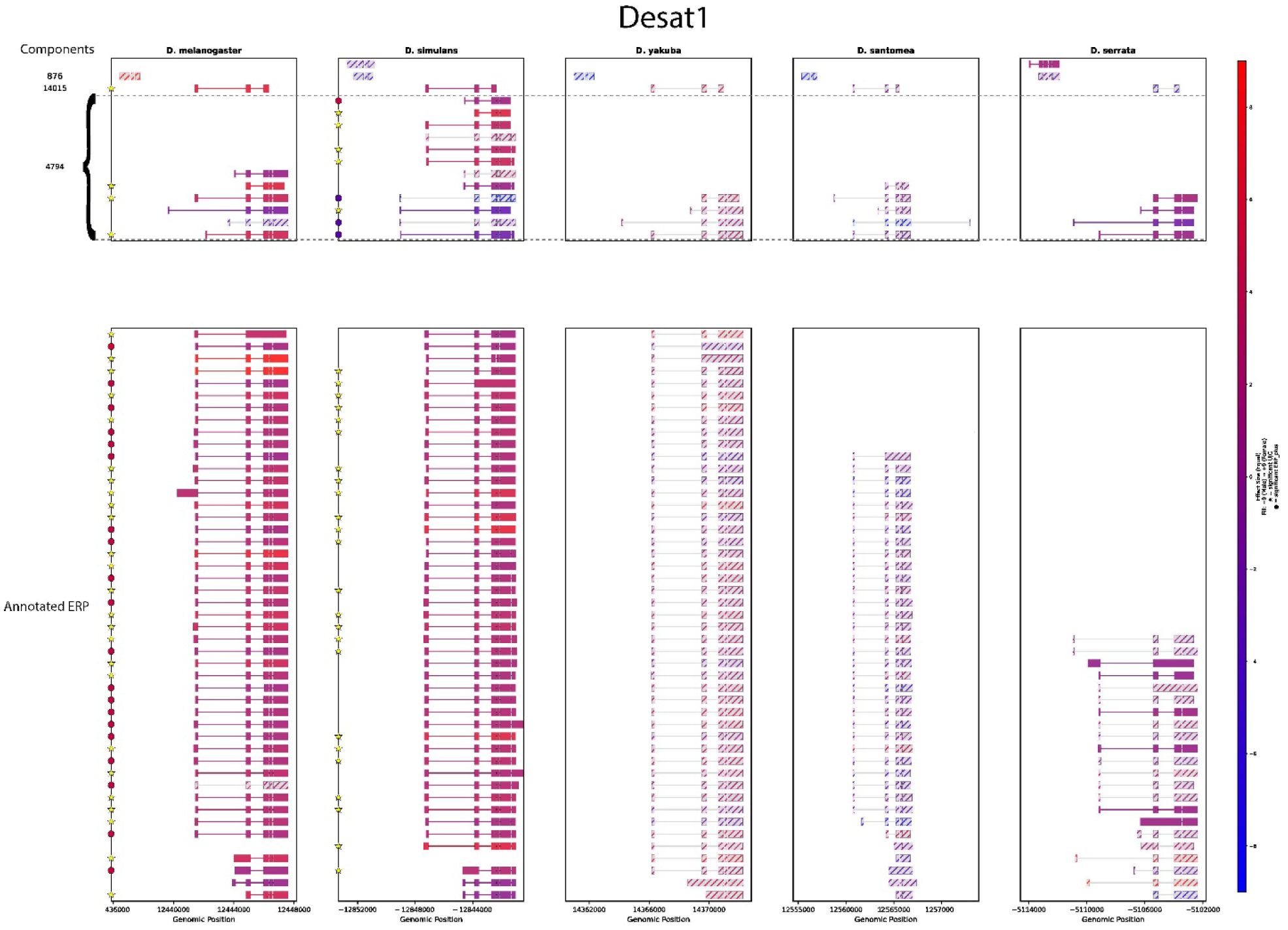
*Desat1* locus. The experimental data for each of the five species. The top row shows annotated transcripts with at least 1 read, the second row shows transcripts with annotated ERP and at least 10 reads. The shading is on a heat map gradient by the effect size (right bar) with red indicating female bias and blue indicating male bias. Statistical significance (p < 0.05) for the transcript is indicated by a yellow star. The filled circle indicates the set of transcripts that share the exon pattern are significant for sex-bias with the shading reflecting the effect size.

